# Functional connectivity drifts during sleep as a marker of fluctuations in the level of consciousness

**DOI:** 10.1101/2025.11.14.688400

**Authors:** João Patriota, Giulia Moreni, Jorge Mejias, Lucia Talamini, Umberto Olcese

## Abstract

During the wake-sleep cycle, consciousness waxes and wanes, and this is thought to be reflected in varying levels of integration between brain areas. Recent studies challenged the notion that consciousness is homogeneously present or absent in a brain state, as exemplified by conscious reports found in otherwise unconscious Non-REM sleep. We tested if functional connectivity between neurons varies within brain states in a way compatible with a fluctuating level of consciousness. We examined directed functional connectivity between neurons across the wake-sleep cycle in rats, at a scale of a few seconds. We analyzed patterns of functional connectivity to determine if Non-REM sleep contains epochs in which inter-areal integration is comparable to that observed in wakefulness and REM sleep, and vice versa. This study will potentially reveal if circuit-level connectivity patterns are observed during sleep stages, in line with the presence of an alternation between levels of consciousness not only between but also within brain states.

## Introduction

Every day consciousness fades when we fall asleep and is restored upon waking up - making sleep a key model for studying the level of consciousness (Nir et al., 2013). However, sleep is not synonymous with a complete absence of consciousness (Koch et al., 2016; Siclari et al., 2017). Early descriptions of rapid eye movement (REM) sleep (Aserinsky & Kleitman, 1953) discussed its association with vivid reports of conscious experiences (dreams). In contrast, Non-REM (NREM) sleep had classically been associated with the polar opposite: oblivion and complete disconnection (Koch et al., 2016) - but see (Siclari et al., 2017). Varying consciousness levels across wakefulness, NREM, and REM sleep may be related to the brain network’s functional integration (Koch et al., 2016; Lu et al., 2012; Massimini et al., 2005; Olcese et al., 2016), as has also been hypothesized by leading theories of consciousness (Storm et al., 2024; Tagliazucchi et al., 2013; Tononi et al., 2024). In line with this, evidence suggests that the network transitions from globally integrated in wakefulness to disintegrated during NREM, but not REM sleep (Massimini et al., 2005; Spoormaker et al., 2012). These studies generally considered sleep stages as homogeneous, and thus quantified the average levels of integration separately per brain state.

However, sleep stages can be highly heterogeneous. First, slow wave activity (SWA) is now understood to be a spatially restricted phenomenon, as indicated by its local nature (Vyazovskiy et al., 2011). Therefore, during NREM sleep different brain regions can exhibit distinct activity profiles. This includes phenomena such as local sleep and wakefulness (D’Ambrosio et al., 2019; Huber et al., 2004; Nobili, 2012; Vyazovskiy et al., 2011). An analogously heterogeneous picture has emerged for what pertains to the presence or absence of consciousness during sleep. For example, dream reports were thought to be exclusive to REM sleep, but consistent accounts of unconscious periods during REM sleep and conscious experiences during NREM sleep challenge this idea (Juan et al., 2023; Siclari et al., 2017). Importantly, variations in EEG oscillatory dynamics within NREM and REM sleep correlate with dream reports, implying a dependency of dreams in local processes. Specifically, dream reports in both NREM and REM were found in a high-density EEG study to correlate with a decrease in slow wave activity and an increase in high-frequency oscillations in the posterior cortex (Siclari et al., 2017). This suggests that, even within brain states traditionally associated with the absence or presence of consciousness (NREM and REM sleep, respectively), spatiotemporally localized patterns of activity can emerge and potentially support conscious mentation.

Moreover, sleep hosts essential processes for physical and mental functions. For instance, sleep’s crucial role in memory consolidation (Rasch & Born, 2013) is reflected in coordinated reactivation of activity between the hippocampus and cortical regions during NREM sleep (Chen & Wilson, 2023; Ji & Wilson, 2007; Sanders et al., 2019), when most of the cortical network appears to be disconnected in comparison to wakefulness (Massimini et al., 2005). Motor learning has long been known to locally modulate the amplitude of slow wave activity during NREM sleep: enhanced low power oscillations are observed in the motor cortex after learning a new motor task (Huber et al., 2004), and the opposite occurs after arm immobilization (Huber et al., 2006). Importantly, a causal link was found between increased slow wave activity and memory consolidation (Marshall et al., 2006). Experience-dependent plasticity has also been shown to enhance coordinated activity between distal areas, as exemplified by the selective synchronous reactivation of memory traces occurring in connected hippocampal and cortical ensembles (Ji & Wilson, 2007; Lansink et al., 2009). Along this line, we previously showed that transitions between brain states do not homogeneously modulate functional connectivity (FC) (Olcese et al., 2016; Olcese, Bos, et al., 2018). Specifically, intra-areal FC is generally preserved between wakefulness and NREM sleep. Conversely, inter-areal FC appears mostly depressed in NREM compared to wakefulness, although communication between hippocampus and cortex is generally preserved, and coupling between neurons significantly modulated during task performance in wakefulness can even be enhanced (Olcese et al., 2016; Olcese, Bos, et al., 2018). However, no study so far - to our knowledge - has investigated the question of whether FC only varies between brain states or also - similar to what occurs for neural activity - also within brain states.

The diversity in the forms of neural activity that has been observed within brain states has led us to hypothesize that the functional architecture present across the brain network might also not be homogeneous within individual brain states. Specifically, while a global decrease in FC during NREM sleep compared to both REM and wakefulness is expected, different types of activity occurring in specific periods of time or subnetworks might lead to distinct levels of integration - and thus consciousness. Here we hypothesize that FC observed in a given time point might deviate in terms of both level of integration and specific connectivity patterns from what is on average observed during a brain state.

We specifically studied whether the FC structure (the set of FC values between individually recorded neurons) within NREM and REM sleep is either homogeneous or not within single brain states. To address this question, we recorded single-unit spiking activity from four brain regions of freely moving rats across wakefulness and sleep. To quantify the FC between individual neurons of different regions across sleep stages we utilized a recently developed method called Current-Based Decomposition (CURBD) (Perich et al., 2020).

Specifically, we hypothesize:

**Hypothesis 1 (H1):** A significant fraction of NREM epochs display a structure of inter-areal FC that is indistinguishable from what is observed on average in REM and/or wakefulness. **Null hypothesis (H0)**: NREM epochs generally exhibit a lower level of FC in comparison to REM and wakefulness or, for those NREM epochs with average FC values comparable to those observed in REM and/or wakefulness, a structure that is different from that observed in REM and wakefulness.

**Hypothesis 2 (H2):** A significant fraction of REM epochs display a structure of inter-areal FC that is indistinguishable from what is observed on average in NREM. **H0**: REM epochs generally exhibit a higher level of FC in comparison to NREM or, for those REM epochs with average FC values comparable to those observed in NREM, a structure that is different from that observed in NREM.

This study investigated sleep-stage-specific neural dynamics, by quantifying how FC changes within and between cortical and subcortical areas across brain states. This deepened our understanding of the nuanced neural dynamics across different sleep stages, potentially shedding light on how such patterns impact cognitive processing and consciousness.

## Methods

In the present study, we utilized a dataset that was originally acquired for a different research project and to answer a separate research question. Specifically, the current study used a portion of the previously collected data that has not yet been used for any analysis. Therefore, the current study will constitute an instance of data re-use. All methods employed to record this dataset will be discussed below, as well as details on how this research question can be answered with this dataset and on considerations about the sample size. Being a data re-use project, we would only be able to reach conclusions if the previously collected dataset provides sufficient statistical power.

The Methods section is structured into five sections. The section “Experimental paradigm” describes the procedures used to collect the data. The section “Analysis plan” focuses on all analytical procedures, from pre-processing to CURBD model fitting. The section “CURBD validation” describes a preliminary analysis that we performed on simulated data to validate the application of the CURBD model and determine the minimal number of neurons for which the approach returns valid results. The section “Statistical plan” describes the hypotheses in depth, how we addressed them and all the necessary statistical analyses that we have planned. Finally, the results of some pilots analysis that we performed to test the proposed methodology are presented in the “Pilot Analysis” section.

### Experimental paradigm

#### Ethical approval

All experiments were performed in accordance with the National Guidelines on Animal Experiments and were approved by the National Animal Experimentation Committee and by the Animal Welfare Committee of the University of Amsterdam. Additionally, we complied with the ARRIVE guidelines (Percie du Sert et al., 2020)

#### General Task Design

Our study employed a within-subject five-day fear conditioning and extinction task. Over the course of five days, we sequentially carried out a series of habituation, fear conditioning, and fear extinction tasks. In this study we utilized a portion of the original dataset, comprising the first three days of experiments (habituation, fear conditioning and fear test). During this period multi-area tetrode recordings were performed across both task performing and rest sessions. The fear conditioning and extinction task is not directly relevant for the purposes of the present study. Specifically, as described more in depth later (section “Experimental procedure”), we expect that any consequence of fear conditioning or extinction on neural activity and connectivity will not affect our research question of whether the FC structure is homogeneous or not within individual brain states. For this reason, we believe that the re-use of an available dataset to answer an independent research question is more ethically responsible compared to performing an ex novo experiment. For the sake of completion, we nonetheless provided details about the complete experimental paradigm that was performed, even when not directly relevant to our research question.

#### Experimental animals

Three male Lister Hooded rats (aged 2-5 months, Envigo, The Netherlands) were housed in pairs during behavioral training. Following the implantation of a tetrode drive, rats were individually housed in transparent cages (53×53×60 cm). All rats were maintained on a regular (i.e., non-reverse) 12-h light/12-h dark schedule and tested in the light (inactive) period. During all experiments rats were fed in an ad libitum regime. After about 5-6 hours of recordings the animals were placed back in their home cage in the animal facility, and allowed to sleep for the rest of the light-on period.

#### Behavioral apparatus

Behavioral measurements were performed in a plexiglass box, with an electric grid (equidistant spaces of 1cm.) on the bottom (Fig 1A; width 53 (base) x length 53 x height 60 cm). Current delivery was controlled by a dedicated scrambled grid current generator (MANHSCK100A, Lafayette). The transparent walls allowed us to present different visual cues via drawings placed on the walls, to further enhance the context of each behavioral stage. Sleep recordings took place in a nest (30 x 30 x 10 cm), placed in the same position in the room as the plexiglass box. Importantly, the nest was only used to allow sleep recordings, but was placed under a different context than the box. During the recordings, auditory stimuli were presented at 60dB (zylux multimedia speaker system, Dell).

**Figure 1.**
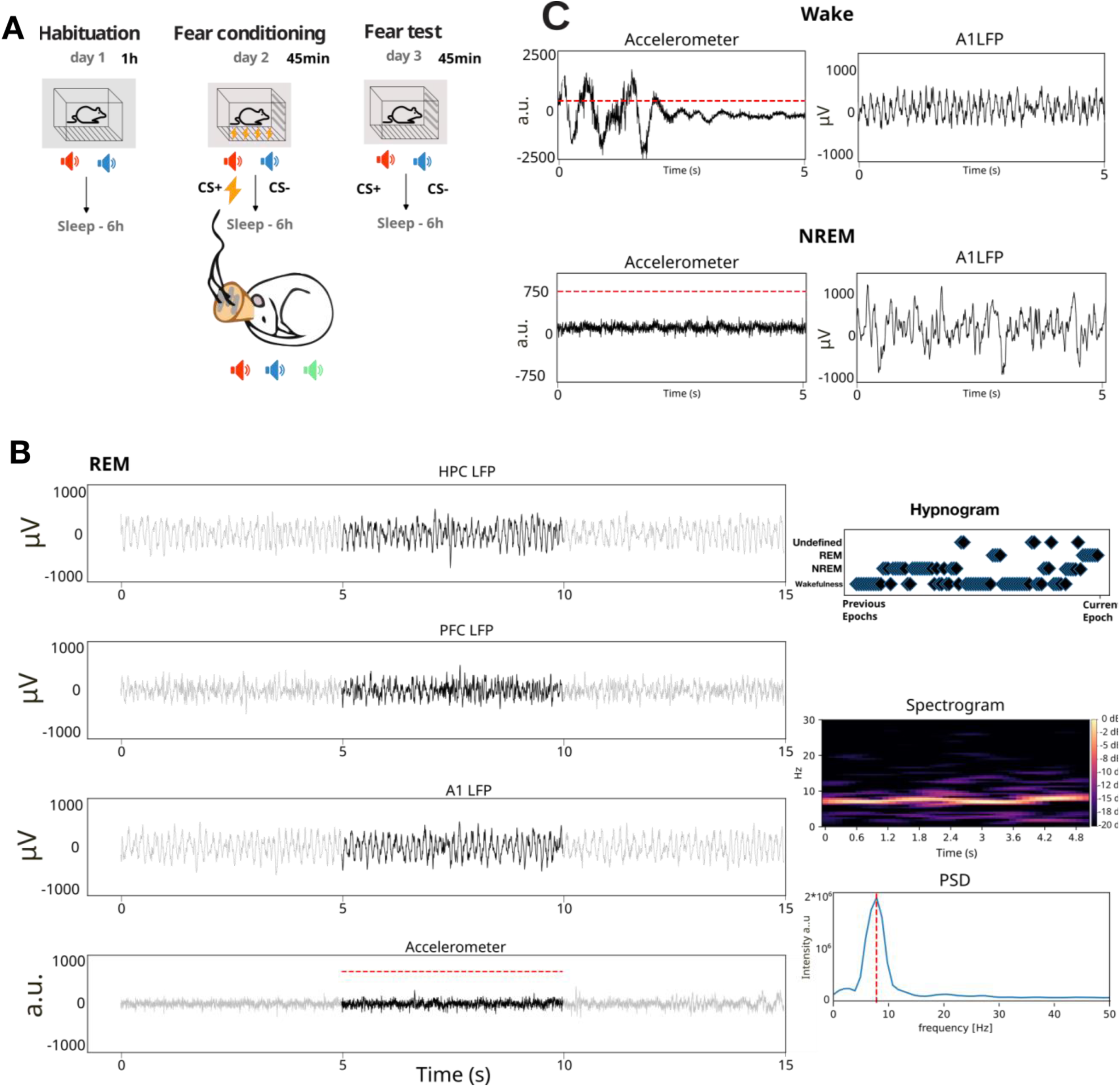
Experimental procedure and scoring of behavioral states. **A**. Day to day experimental procedure. Each experimental session (Habituation, Fear conditioning and Fear test) was followed by a session of sleep recordings. **B**. Exemplary traces of REM classification, indicating all the information that was used to perform sleep stage classification. In the left column (from top to bottom) LFP activity is depicted for the different channels (HPC, PFC, A1) that were used to monitor global ongoing activity. Gray traces indicate previous and subsequent sleep epoch, and the black trace is the epoch we are scoring. Below the LFP traces, accelerometer activity is depicted, with the dashed red line indicating the defined threshold for motor activity. On the right, from top to bottom: **Hypnogram** scoring of the previous 100 epochs; **spectrogram** showing the time frequency decomposition for the current epoch based on cortical (A1) activity (note that the oscillatory activity is centered on 8 Hz); **Power source density** (PSD) of the current epoch, indicating that most power is within the theta (6 - 12 Hz) band. **C**. Example LFP traces from one cortical channel (A1) recorded, respectively, during WAKE and NREM. Note the predominance of lower frequencies and higher amplitude during NREM, as well as the lower levels of head acceleration.

#### Behavioral paradigm

In preparation for the experiment, animals were gradually acclimatized to the experimental setup over a two-week period. Each animal was familiarized daily with the experimental box by spending at least thirty minutes within it, followed by an equal amount of time spent in the nest (Fig 1A).

Upon completion of this acclimatization phase, electrode implantation followed. The behavioral paradigm was initiated at least fourteen days after surgery, to ensure full health of the animal, and lasted for up to five consecutive days. Each daily experimental session was divided into two distinct phases. In the first phase, during about 60 minutes, animals were given freedom to navigate the experimental box while engaging in the assigned task. Once this task was completed, the subjects were relocated to the nest for sleep recordings that took place for about 5h.

Detailed descriptions of each experimental day’s activities are elaborated upon in the subsequent section.

#### Detailed Experimental Procedure

In this experiment, we followed an auditory cue fear conditioning paradigm (Bagur et al., 2021; LeDoux, 2014).

**Day 1: Habituation** - The initial day of the experiment entailed habituation. Each animal was placed in the experimental box and we initiated the habituation phase. This phase involved 25 presentations of two types of auditory stimuli (Conditioned Stimuli or CS, 2s white noise filtered at 3kHz, and white noise filtered at 9kHz, respectively). Each CS presentation was interspersed with a randomized inter-trial interval (ITI) of 3 to 5 minutes. The auditory stimuli were presented in a randomized sequence.

**Day 2: Fear Conditioning** - On the second day, each animal was placed in the experimental box. Visual cues were added to enrich the context (Fig 1A). Animals were then subjected to fear conditioning procedures. Post baseline period (3 minutes), the animals were introduced to 10 trials comprising five CS− and five paired CS+-Unconditioned Stimulus (US) presentations (Bagur et al., 2021). The same CS stimuli used in day one were presented, with one stimulus being used as CS+, and the other as CS−. The US, a foot shock (500ms, 0.5mA), was delivered immediately following the end of the sound. The ITI was randomized between 3 to 5 minutes. Each sound lasted 2 seconds, with the shock administered instantly after the sound via a specified controller.

**Day 3: Fear Test** - On the third day, the probe test was conducted. Animals were placed in the experimental box, visual aid on the plexiglass walls were the same used during the fear conditioning step. Following the baseline period (3 minutes), the animals were presented with 12 trials including five CS− and seven CS+ pairs (Bagur et al., 2021). Sound presentation was conducted in a block format: initially, all CS+ sounds were presented, followed by a block of CS−sounds. The ITI was again randomized between 3 to 5 minutes, with each sound lasting 2 seconds.

Subsequent experimental procedures, unrelated to the primary focus of this study, spanned another two days, after which the animal was euthanized.

**Day 4: Extinction training -** On the fourth day, each animal was placed in the experimental box and we initiated the extinction phase. Here the visual cues were different from the ones used during days two and three. This phase involved 30 presentations of both CS+ and CS−. Each CS presentation was interspersed with a randomized inter-trial interval (ITI) of 3 to 5 minutes. The auditory stimuli were presented in a randomized sequence.

**Day 5: Extinction test** - On the fifth day, the extinction test was conducted. Following the baseline period, the animals were presented with 12 trials including five CS− and seven CS+ pairs (Bagur et al., 2021). Sound presentation was conducted in a block format: initially, all CS+ sounds were presented, followed by a block of CS− sounds. The ITI was again randomized between 3 to 5 minutes, with each sound lasting 2 seconds.

Here we combined data recorded across different experimental conditions, under the assumption that brain states will similarly modulate inter-areal communication. In other words, we expected that the homogeneous or heterogeneous nature of the FC structure within individual brain states would not depend on the experimental phase. However, it is likely that each experimental condition might differentially affect how brain states modulated functional coupling between brain regions (i.e., the details of the FC structure itself). Previous literature suggests that this effect would be heterogeneous, and would differentially impact not only different brain areas, but also specific neuronal subpopulations within areas - see e.g.(Ji & Wilson, 2007; Lansink et al., 2009; Olcese et al., 2016). Based on the results that we obtained, this might be a topic for further investigation.

**Sleep Recordings** - Sleep sessions were conducted immediately after each behavioral session. Animals were relocated to a dimly lit flower pot for sleep recordings, each session lasting approximately five hours. Upon identifying sleep onset, a period defined by the emergence of significant slow-wave activity (SWA) in the cortical signals, coupled with a notable decrease in the animal’s motion (see also later), as detected via our camera-based monitoring system, both CS+ and CS− stimuli were randomly played, supplemented by a novel sound (white noise filtered at 6kHz, 2s). Inter stimulus interval was randomly set between 2 and 4 seconds. If an arousal was observed by the experimenter, sound presentation ceased immediately. Arousals were identified as any significant disruptions in ongoing stereotypical wave patterns, such as theta waves during REM sleep or delta waves during NREM sleep. Additionally, these disruptions were concurrently verified through motion observations, providing a multimodal approach to effectively detect instances of arousal in the animals. Each session featured approximately 200 sound presentations.

#### Electrode implantation and surgery

In order to allow full recovery and to avoid interferences on the experimental procedures, surgery was performed at least 14 days prior to the first experimental day (habituation). Tetrodes were constructed from four twisted 13-µm coated nichrome wires (California Fine Wire). The electrode tips were gold-plated to reduce electrode impedances to 300-800 kΩ at 1 kHz. Tetrodes were loaded into a custom-built microdrive, containing 36 individually movable tetrodes (Bos et al., 2017; Lansink et al., 2007). The drives targeted eight tetrodes to the primary auditory cortex (A1; target coordinates in mm: −4.81 AP, −3.81 ML, 4.32 DV (Paxinos and Watson, 2007)), eight tetrodes to dorsal hippocampal CA1 area (−3.2 AP, −1.51 ML, 2.88 DV), eight tetrodes to the basolateral amygdala (BLA; −2.74 AP, −3.35 ML, 7.2 DV) and eight tetrodes to the prefrontal cortex (mPFC; +1.83 AP, −0.83 ML, 5.33 DV).

Prior to surgery, rats received a subcutaneous injection of buprenorphine (Buprecare, 0.04 mg kg-1), meloxicam (Metacam, 2 mg kg-1) and enrofloxacin (Baytril, 5 mg kg-1). Anesthesia was induced by placing the animal in a closed plexiglass box filled with isoflurane vapor (3.0%). During surgery the animals were mounted in a stereotaxic frame where anesthesia was maintained using isoflurane (1.0-2.0%) and body temperature was maintained between 35 and 36 deg using a heating pad. Local anesthetic (lidocaine) was applied directly on the periosteum before exposing the skull. The skull was thoroughly cleaned with a 3% hydrogen peroxide solution to roughen the skull surface. The hydrogen peroxide was removed by rinsing the skull three times using saline. Six screws, from which one in the occipital bone served as ground, were inserted into the skull to improve the stability of the implant. After craniotomy and durotomy were performed, the drive was positioned over the craniotomies. The craniotomies were sealed using silicone adhesive (Kwik-Sil), the skull was covered with a layer of dental adhesive (OptiBond, Kerr) and primer (3M Transbond) and finally the drive was anchored using dental cement (Kemdent). Finally, all tetrodes were lowered into the superficial layers of the cortex. From the third day, tetrodes were gradually lowered towards their target location regions over a span of 7-14 days. Post-operative care included a subcutaneous injection of meloxicam once per day for two days following surgery, and a single injection of enrofloxacin on the day following surgery.

#### Data acquisition

For data collection, the rat was connected to the recording equipment (Open Ephys, 148 channels) via operational amplifiers (RHD2000, Intan) and a tether cable connected to a commutator (Saturn-5, Neuralynx). After the post-surgery recovery period, tetrodes were lowered to their target location while their current location in the brain was estimated based on the distance the tetrode was moved in the drive and through thorough assessment of the LFP and spike signals. In order to sample activity from a diverse population of neurons, tetrodes were independently advanced after each recording session by at least 250 um. Each tetrode was moved independently to a location where spiking activity was clearly visible in the raw trace. During experiments the neural activity was acquired at 30 kHz continuously and stored on a hard drive for further processing. A camera (acA1920-25gm, Basler) positioned above the behavioral setup recorded behavioral activity. The behavioral apparatus was controlled using an Arduino Mega (Arduino). Programmed commands and events recorded were sent to the Open Ephys recording system as TTL pulses. Animals were equipped with a headstage (Intan RHD2132 board) which includes a 3-axis accelerometer that gives access to the animals’ acceleration with high temporal resolution.

#### Histology

After the final recording session, currents (12 µA, 10 s) were applied to one lead of each tetrode to mark its endpoint with a small lesion. Twenty-four hours after lesioning, the animals were deeply anesthetized with Pentobarbital (Nembutal, Ceva Sante Animale, 60 mg ml-1, 1.0 ml intraperitoneal) and transcardially perfused with saline followed by a 4% paraformaldehyde solution (pH 7.4, phosphate buffered). After a minimum of 24h post-fixation, the brain was sectioned into 40-50µm thick slices using a vibratome. A Nissl staining (Cresyl Violet) was set to mark cell bodies and to reconstruct tetrode tracks and localize their endpoints. The sections were imaged and aligned to the 3D Waxholm reference atlas (Papp et al., 2014)

### Analysis Plan

#### Motion tracking

Animal recordings were equipped with a headstage (Intan RHD2132 board) which included a 3-axis accelerometer which gave access to the animal’s acceleration with high temporal resolution. This information was used to measure motor activity in the same temporal resolution as the collected local field potential activity (Bagur et al., 2021).

#### Spike detection

Signals were digitally band-pass filtered between 800 and 6000 Hz for spike recordings. Action potentials were assigned to single neurons by using Tridesclous, a semiautomated spike sorting algorithm (Garcia & Pouzat, 2019) implemented within the Python package spikeinterface (Buccino et al., 2020), and manually curated using the Phy GUI (Rossant et al., 2016). Each candidate unit was considered for further inspection during manual curation based on its waveform and autocorrelation function. High-quality single units were included, and defined as having (1) an isolation distance higher than 10, (2) less than 0.1% of their spikes within the refractory period of 1.5 ms, (3) spiking activity present throughout the session. Other candidate units, with physiological waveforms but with more than 0.1% of their spikes within the refractory period (i.e., still respecting criteria number 1 and 3), were deemed as high-quality multi units and were included for the CURBD analysis. Local field potentials (LFPs) were recorded from all tetrodes, continuously sampled at 30 kHz, and band-pass filtered between 1 and 1000 Hz. Across 7 sessions, we obtained on average, per area (mean ± standard deviation): 6.3 ± 1.4 in BLA, 6.6 ± 3.1 in PFC, 5.9 ± 4.64 in A1, 3.43 ± 1.29 in HPC (Table 5). For further description of the dataset, see the sampling plan below.

#### Sleep scoring

Considering the fragmented nature of rodent sleep, we segmented the recording sessions into discrete 5-second epochs (Olcese, Bos, et al., 2018), individually scored for analysis (Fig 1B). This scoring was done by simultaneously evaluating several factors: cortical Local Field Potential (LFP) channels, LFP power spectrum within each epoch and motor activity (ascertained through accelerometer displacement). More in detail, sleep stage scoring was done by taking into account LFP oscillatory dynamics in each epoch of multiple cortical recording channels as well as the level of motor activity (measured in terms of head acceleration). Epochs that displayed 1) low levels of cortical slow-wave activity (total power between 0.5 and 4 Hz) and detectable levels of motor activity (head acceleration over a visually determined threshold of 750 arbitrary units - see Fig 1C) or 2) no detectable motor activity in said epoch, but with motor activity above the threshold in the previous 10s (two epochs) or subsequent 10s were classified as wakefulness (WAKE). The motor threshold was determined by measuring head acceleration during periods when video recordings showed freezing or immobility in the absence of motor actions such as grooming (dos Santos Lima et al., 2019). We also classified an epoch as WAKE if, on top of the lack of movement, we did not observe theta oscillations. The presence of theta oscillations (in the absence of motion) was instead taken as an indication of REM sleep. Epochs with a high level of slow-wave activity and limited motion were scored as NREM. Overall, this evaluation allowed us to classify the behavioral state of recorded animals into three distinct stages: wakefulness (WAKE), NREM sleep and REM sleep. In case the typical markers of individual sleep stages were absent or were not consistently present in all recorded areas, we classified such epochs as undefined (UND). This was for example the case in the transition between sleep stages, when epochs can show intermediate patterns of activity either temporally (within each epoch) or spatially (across areas) (Emrick et al., 2016). Such epochs were excluded from further analysis. Importantly, this exclusion criterion makes it unlikely that, even in the absence of EEG and EMG recordings, we will spuriously include local brain states (Vyazovskiy et al., 2011) in our analysis. Specifically, our recording sites span from posterior areas (auditory cortex) to frontal ones (prefrontal cortex) and even deep regions (hippocampus, basolateral amygdala). Therefore, we deem it very unlikely that any (region-specific) sleep stage marker simultaneously detected in all recording sites would reflect a local rather than global phenomenon.

These stringent criteria, which include a conservative acceleration threshold to detect immobility (Bagur et al., 2021; dos Santos Lima et al., 2019), were used to identify motor activity and to facilitate the identification of sleep stages, contributing to the validity of our results. We corroborated the outcomes of our behavioral scoring procedure by separately plotting several electrophysiological and behavioral parameters for each recording session (Fig 1B). Hippocampal ripples, typical of quiet wakefulness and NREM but less frequent in active wakefulness, further validated the effectiveness of our scoring procedure.

Across 7 sessions we obtained, on average, per sleep stage: 2023.86 ± 523.74 NREM epochs, 779.57 ± 224.64 REM epochs and 1242.43 ± 445.31 epochs of wakefulness. For further description of the dataset, see the sampling plan below. Epochs that failed to meet the criteria for any of these four states were discarded (Olcese et al., 2016; Olcese, Bos, et al., 2018).

We then employed a custom script to generate comprehensive information on time-frequency, power source distribution, downsampled data visualization, and animal motion. This process provided a platform to visualize the data, plot 5-second epochs, and subsequently calculate the Hilbert amplitudes (smoothed with a Gaussian kernel, σ = 1 s) of filtered theta (6–12 Hz) and delta (1–4 Hz) band LFP (Fig 1B).

#### Estimation of connectivity

Building upon the already done preprocessing (spike sorting and sleep stage categorization), we separated Single Unit Activity (SUA) into distinct trials for each 5 s sleep epoch. Current-Based Decomposition (CURBD) (Perich et al., 2020) was applied on an epoch by epoch basis to the spiking trains from different pairs of neurons belonging to different regions, in order to obtain the connectivity magnitude of all combinations per each epoch. Compared to common methods to study interactions between brain regions such as linear regression, canonical correlation analysis (CCA), constrained dimensionality reduction, generalized linear models (GLMs), or Granger causality, amongst others, CURBD aims to reconstruct underlying connectivity by reproducing the observed activity patterns, rather than just computing (directional) correlations between activity patterns. In this way, CURBD allows us to, in an unbiased way, capture the directionality and magnitude of the interactions within and across regions that can be interpreted as being computationally responsible for the observed neural dynamics (Fig 4A). Among other strategies to fit recurrent neural networks to ensemble recordings - see e.g. (Das & Fiete, 2020; Finkelstein et al., 2021; Kuan et al., 2024; Pandarinath et al., 2018) - CURBD stands out as it was specifically developed to infer functional connectivity between brain areas. This approach is furthermore highly valuable for our purposes, since it is valid even when applied to a low number of units (Perich et al., 2020). Importantly, this method jointly captures mono- and polysynaptic connectivity. This method thus allows to jointly quantify both direct and indirect connections between pairs of regions, providing indications about the overall strength and patterns of interareal connectivity. Since both direct and indirect connections between - but not within - brain regions are thought to be strongly affected by brain states (El-Baba et al., 2019; Jobst et al., 2017; Olcese et al., 2016; Olcese, Bos, et al., 2018), this method is ideally suited to determine to what extent individual brain states are homogenous in terms of multi-regional functional connectivity structures.

CURBD assumes that the high degree of recurrent connectivity present between brain regions makes a recurrent neural network (RNN) suitable to reproduce such dynamics, and that the activity that drives a unit in an RNN can be computed as a weighted sum of the activity of all other source units in the network. CURBD then fits a single layer RNN to neural activity obtained from an experiment, estimating functional connectivity with a high spatiotemporal resolution, taking in account each epoch and at the level of single neurons, but also constrained by the particularities of the experimental data in view of the multi-regional nature of our recordings. A model RNN is constructed such that each unit is trained to match a single experimentally recorded epoch from a neuron from the full dataset of neural population activity from multiple interacting regions. Importantly, the output obtained by CURBD should be considered a model of the data itself - an in silico representation of the experiment: such a model does not incorporate prior assumptions regarding the identity of the model neurons, nor does it consider anatomical constraints. Training occurs where the connectivity matrix of the model RNN is modified over time until the activity of the RNN units match the experimental data (Perich et al., 2020). For each pair of neurons per epoch we obtained a 5 s time series where each point corresponds to the estimated connectivity between those two neurons in that epoch. We then averaged the pairwise connectivity values obtained over the whole 5 s epoch, for all pairs of neurons located in different brain regions, in order to obtain one single pairwise connectivity value per epoch (Fig 4B). Therefore, in the following analyses, we did not consider how connectivity between individual pairs of neurons varies within and between brain states, but rather how the average value of pairwise interactions between areas is modulated. Thus, we did not aim to reconstruct the connectivity diagram between the recorded neurons, but only focus on the average connection strength between regions. While connectivity was estimated independently for each 5 s epoch, results were analyzed first at the level of each recording session, and then combined across sessions. Details are provided in the following sections.

As discussed, CURBD was designed to infer interactions in experimental datasets. However, the method has not been yet validated in a peer-review publication. Therefore, we proceeded to perform a pilot analysis to determine whether CURBD is capable of capturing the dynamics of interareal functional connectivity in a biologically inspired spiking neural model based on previous work (Joglekar et al., 2018; Mejias & Longtin, 2014; Moreni et al., 2023, 2024; Nunes et al., 2021; Olcese et al., 2010). Basically, this toy model allowed us to simulate realistic spiking patterns in a 4-region cortical network in which we could set, on the basis of experimental data (Oh et al., 2014), the average level of monosynaptic connectivity between areas, and also perform current injections in one area and simulate how evoked spiking activity differentially propagates to other areas based on the underlying network connectivity. Importantly, the model was not designed to reproduce the activity of the areas from which recordings were performed. Rather, it only served as a toy model to validate the functionality of CURBD, besides what has been presented in the manuscript proposing the method. The next sections describe the model in depth.

### CURBD validation

#### Model architecture

The computational model, whose architecture is depicted in Fig 2A, is composed of a total number (N_total_) of 5,000 neurons. The model consists of four areas (R1-R4), each containing 85% excitatory neurons (E) and 15% inhibitory (I) neurons (Billeh et al., 2020). Each area thus contains 1062 excitatory cells and 188 inhibitory cells. All neurons receive background noise from the rest of the brain.

**Figure 2.**
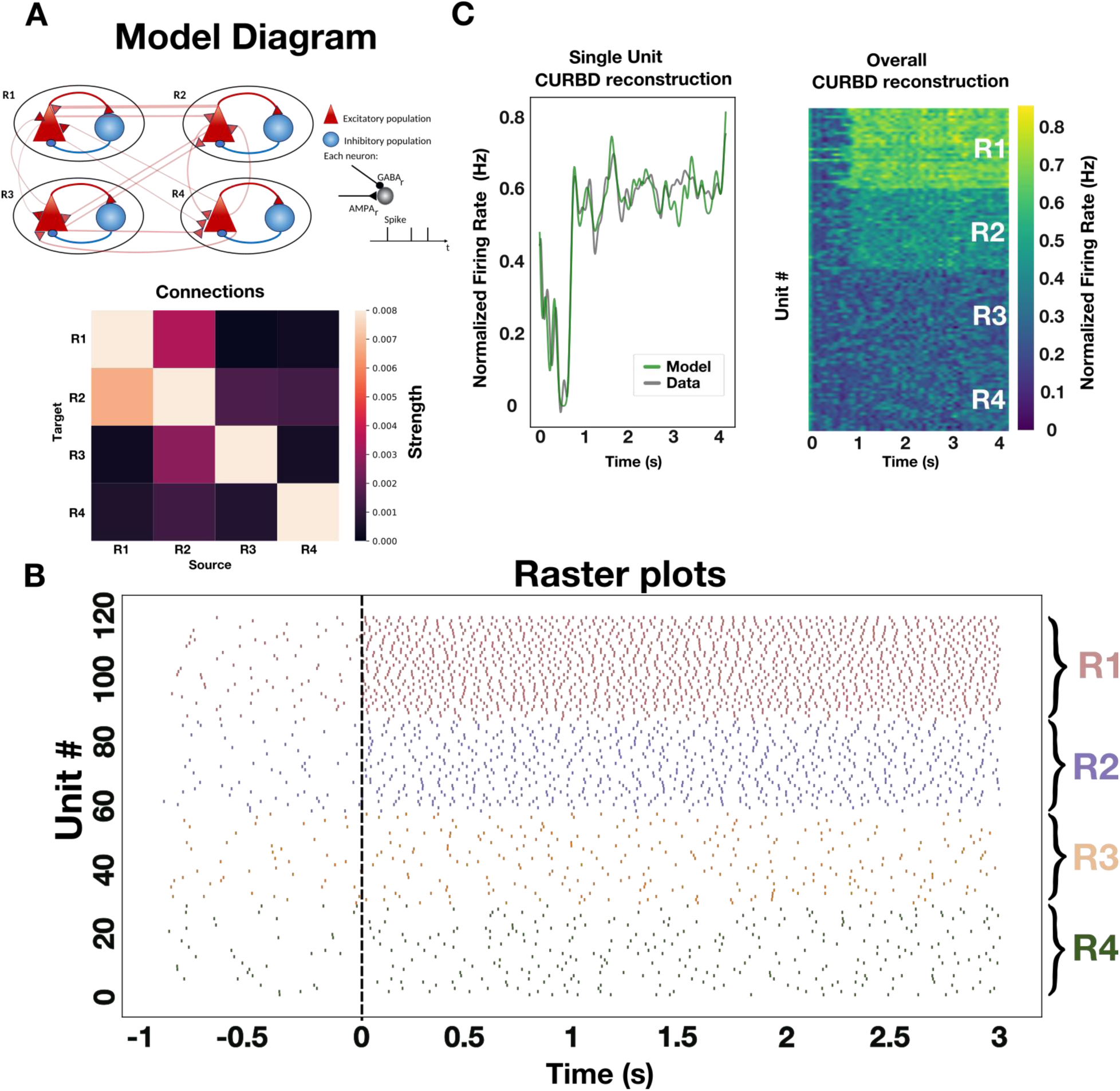
Applying CURBD to a toy model. **A.** Schematics of the model architecture. The model is composed of four areas (R1-R4). In each area one excitatory population (Red triangle) and one inhibitory population (blue circle) are present. In each area 85% of the population is of excitatory neurons and 15% of inhibitory neurons. In total, we simulated 5000 neurons. The thickness of the connections between the symbols represent the model connectivity strength. The heatmap shows the normalized connectivity strength (normalized number of synaptic connections) between each pair of regions. **B.** Raster Plot of spiking activity simulated for 4s. At time 1, a current of 30pA is applied to region 1 (R1). Activity in region 2 (R2), reflecting the stronger connectivity region R1, shows a higher firing rate as well following current injection, indicating the propagation of information between both areas.. **C**. Left. Following training, the CURBD reconstruction (green) of the activity of a single neuron reproduces the activity of the simulated neuron (gray). Activity is expressed as peristimulus time histogram (PSTH) traces. Right. CURBD reconstruction of the activity of all units provided to the algorithm (20 neurons/area). Note that, following current injection, the units in R1 and R2 exhibit higher firing rates, similar to what is observed in the simulated data.

### Single neurons

All excitatory and inhibitory cells in the network are modeled as leaky integrate-and-fire neurons. Each of the two types of cell is characterized by its own set of parameters: a resting potential V_rest_, a firing threshold V_th_, a membrane capacitance C_m_, a membrane leak conductance g_L_ and a refractory period t_ref_. The corresponding membrane time constant is t_m_= C_m_/g_L_. The membrane potential V(t) of a cell is given by:

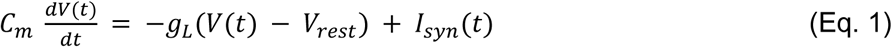

where I_syn_(t) represents the total synaptic current flowing in the cell.

At each time point of simulation, a neuron integrates the total incoming current I_syn_(t) to update its membrane potential V(t). When the threshold V_th_ is reached a spike is generated, followed by an instantaneous reset of the membrane potential to the resting membrane potential V_rest_. Then, for a refractory period t_ref_, the membrane potential stays at its resting value V_rest_ and no spikes can be generated. After t_ref_ has passed, the membrane potential can be updated again (see Table 1 for the corresponding parameter values).

**Table 1:** Parameters values for the two groups of cells. The values are taken from the Allen databas (https://portal.brain-map.org/explore/models/mv1-all-layers) of cells in layer 2/3 of V1, as an example of a realistic network and without loss of generality of our method.

|  | E | I |
| --- | --- | --- |
| $C_m$ (pF) | 123.41 | 70.95 |
| $g_L$ (nS) | 2.47 | 9.49 |
| $t_{ref}$ (ms) | 3 | 1.26 |
| $V_{rest}$ (mV) | -80.97 | -82.35 |
| $V_{th}$ (mV) | -40.53 | -56.32 |

### Parameters of neuron models

Each type of cell (I or E) is characterized by its own set of parameters: a resting potential V_rest_, a firing threshold V_th_, a membrane capacitance C_m_, a membrane leak conductance g_L_ and a refractory period t_ref_.

These data are taken from the Allen institute database (https://portal.brain-map.org/explore/models/mv1-all-layers) of layer 2/3 of the primary visual cortex (V1), reflecting the fact that electrophysiological properties of neurons in that area have been thoroughly studied. Without loss of generality, we chose layer 2/3 as our representative subcircuit due to its particularly strong recurrent excitatory connectivity, which allowed us to design a model with a clear segregation of neurons into separate regions. We use data from pyramidal cells to model our excitatory neurons and data from PV interneurons to model our inhibitory neurons.

### Synapses

The inputs to model neurons consist of three main components: background noise, external (e.g. sensory) input and recurrent input from the modeled areas. EPSCs due to background noise are mediated in the model by AMPA receptors (I_ext,AMPA_(t)) and EPSCs due to external stimuli (i.e. originating from outside the area) are represented by I_ext_(t). The recurrent input from within the model is given by the sum of I_AMPA_(t) and I_GABA_(t). These are all the inputs from all the other presynaptic neurons projecting to the neuron under consideration.

The total synaptic current that each neuron receives is given by:

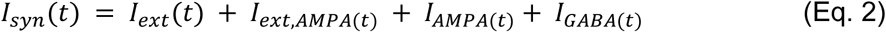

with the first term on the right hand (I_ext_) side being the current injected externally (a constant current of 30 pA whenever the network receives external input, whose value was empirically determined to achieve regular model dynamics), and with the last three terms given by

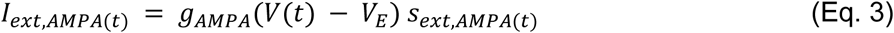

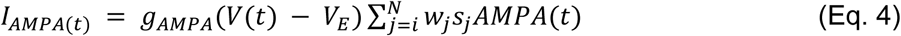

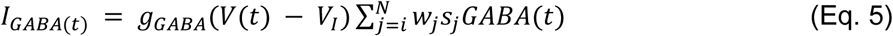

where the reversal potentials are V_E_=0 mV, V_I_ = V_rest_. The g terms represent the conductances of the specific receptor types, and are equal to one for simplicity. The weights w_j_ represent the strength of each synapse received by the neuron. The sum runs over all presynaptic neurons j projecting to the neuron under consideration. The s terms represent the gating variables, or fraction of open channels and their behavior is governed by the following equations

The AMPAR channels are described by

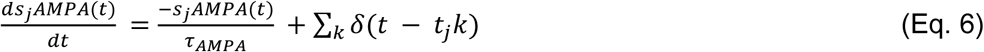

where the time constant of the AMPA currents is t_AMPA_= 2 ms (Wang, 2002) - see also (Hestrin et al., 1990; Spruston et al., 1995) - and the sum over k represents the contribution of all spikes (indicated by delta, δ) emitted by presynaptic neuron j. In the case of external AMPA currents (Eq. 3), the spikes are emitted accordingly to a Poisson process with rate u_bkgnd_. The two groups of cells in each area (I and E) are receiving a different Poisson rate of background noise: u_bkgnd_ = 930 Hz for excitatory cells and u_bkgnd_ = 1460 Hz for inhibitory cells. These values were empirically determined, in line with what is commonly done for similar models (Potjans & Diesmann, 2014; Van Albada et al., 2015). This is roughly equivalent to having roughly ∼1000 external neurons projecting to each neuron in either group and firing at around 1 and 1.5 spikes/s respectively, which constitutes a good approximation for spontaneous activity levels in cortex

The GABA_A_ receptor synaptic variable is described by

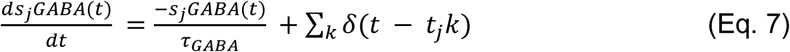

where the time constant of GABA_A_ receptor current is 5 ms (Wang, 2002) - see also (Salin & Prince, 1996; Xiang et al., 1998).

### Connectivity and synaptic weights

Each neuron in the network projects a synapse to any other neuron with a certain probability which depends on the cell group (i.e. excitatory or inhibitory) and area (i.e. R1, R2…). The values in matrix P (Table 2) indicate the probability that a neuron in group A (e.g. an inhibitory cell in area R1) is connected to a neuron in group B (e.g. excitatory cell in area R1). The weight of each existing synapse w_j_ from neurons of group A to neurons in group B in the same area is chosen to be equal to

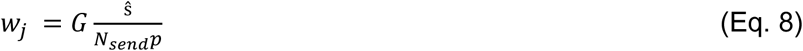

where G=5 is the global coupling factor (this value was empirically chosen to guarantee a minimum level of strength of the overall connectivity without leading to pathological synchrony in the dynamics), S is the overall strength between the two connected groups of cells, N_send_ is the number of neurons in the sending population A, and p the probability of connection between the neurons of the two groups (A and B) taken from the connection probability matrix (Table 2), S is from the experimental synaptic connectivity matrix (Table 3). As for the other model parameters, the values are chosen from layer 2/3 of the cortical column in V1 (Billeh et al., 2020). Complete matrices for the values of S and p of the cortical column where these values were taken from can be found at https://portal.brain-map.org/explore/models/mv1-all-layers. The normalization in Eq. 8 above guarantees that the dynamics and equilibrium points of the system scale properly with the size of the network.

**Table 2:** Within-area connection probability. Probability of connections between populations in the same area. Sending population is indicated in the header of the first column of the table and receiving population is indicated in the header of the first row of the table. All values were taken from (Billeh et al., 2020).

|  |  | Target |  |
| --- | --- | --- | --- |
|  |  | E | I |
| Source | E | 0.16 | 0.395 |
|  | I | 0.411 | 0.451 |

**Table 3:** Connection strength within area. Same as Table 2, but for intra-area connection strength S.

|  |  | Target |  |
| --- | --- | --- | --- |
|  |  | E | I |
| Source | E | 0.36 | 0.149 |
|  | I | 0.48 | 0.68 |

Here we use the term connection with reference to subpopulations or groups, defined by the pre- and postsynaptic neuron types in each area (I and E cells). The connection probability defines the probability for each possible pair of pre- and postsynaptic neurons to form a connection between them. If p=0.1 this connects all neuron pairs of the two groups with a probability of 10%. The connectivity probability matrix P within one area is defined by the 2×2 = 4 connection probabilities p between the 2 considered cell groups (E and I cells). For simplicity, and without loss of generality for our method, the same values are used for all the 4 areas. Then the four areas are connected only through excitatory neurons (see Fig 2A) with a fixed probability p=0.3 (empirically determined).

Each connection also has a particular strength which differs per neuron group. Thus, the strength was specified at the level of neuron type X projecting to neuron type Y (Table 3). These values are rescaled to find the single synaptic strength w_j_ between neurons to use in the model (Eq. 8).

As mentioned earlier, the four areas R1-R4 are connected only through pyramidal cells (See Fig 2A). The connections strength between excitatory cells (E) of the four different areas are taken from experimental data (Oh et al., 2014) and are the values between four brain areas (Primary visual area, Anterolateral visual area, Barrel field in the primary somatosensory area, Dorsal part of the anterior cingulate area) which are represented by area R1-R4 in our model. We chose these areas to have a spatially distributed network including both sensory and association areas, but other areas can be easily considered as well. The values of the connection strength of single synapse w_j_ are reported in the following table (Table 4). Note how the connectivity between R1 and R2 is one order of magnitude higher than that between all other areas, to test if CURBD can capture this enhanced connectivity. These values are then increased by a global constant G=50, selected empirically to obtain realistic firing rates in all 4 areas.

**Table 4:**
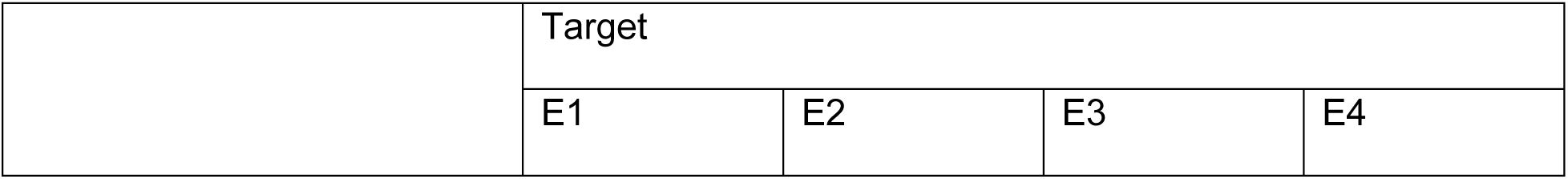
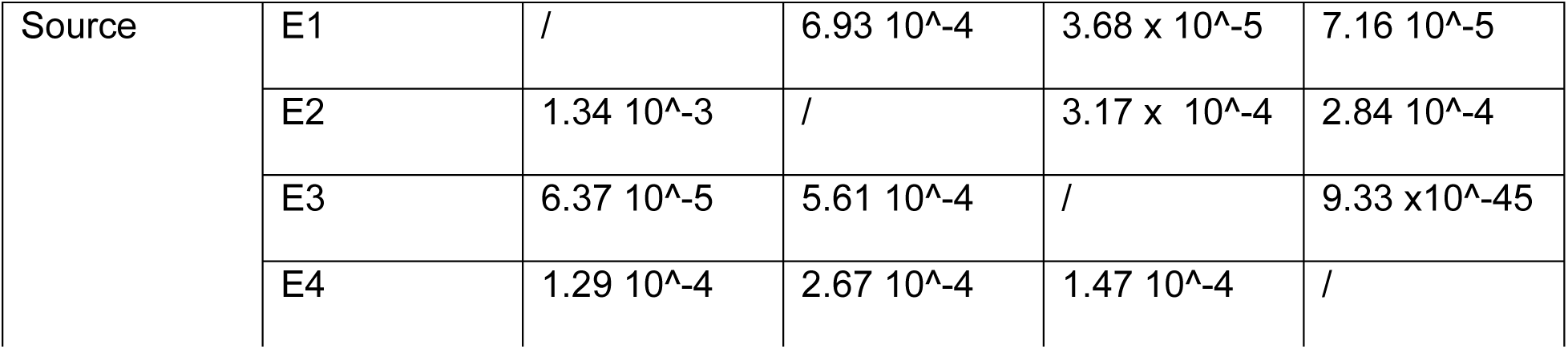
Synaptic strength between excitatory neurons in the four different areas (E1: excitatory neurons in R1, etc.).

|  | Target |  |  |  |
| --- | --- | --- | --- | --- |
|  | E1 | E2 | E3 | E4 |

|  |  |  |  |  |  |
| --- | --- | --- | --- | --- | --- |
| Source | E1 | / | $6.93 \times 10^{-4}$ | $3.68 \times 10^{-5}$ | $7.16 \times 10^{-5}$ |
| | E2 | $1.34 \times 10^{-3}$ | / | $3.17 \times 10^{-4}$ | $2.84 \times 10^{-4}$ |
| | E3 | $6.37 \times 10^{-5}$ | $5.61 \times 10^{-4}$ | / | $9.33 \times 10^{-45}$ |
| | E4 | $1.29 \times 10^{-4}$ | $2.67 \times 10^{-4}$ | $1.47 \times 10^{-4}$ | / |

### Simulation details

All simulations were performed using custom Python code and Brian 2.0. Differential equations were solved using an Euler-Mayurama algorithm with an integration step of 0.1 ms. The code will be made available via GitHub upon acceptance of the manuscript.

#### CURBD validation on simulated data

The chosen model implementation allowed us to explore if CURBD could, on the basis of the activity patterns observed in different regions, capture the predetermined inte-areal connectivity values. In this implementation, we selected connection strength between regions that are similar in magnitude from what is obtained via experimental data (Oh et al., 2014). Here, the pair R1-R2 is more strongly connected than other pairs such as R3-R4, R1-R3, etc. (Fig 2A), so we expect that, if CURBD can correctly determine the functional connectivity levels between all pairs of regions, it should assign the highest levels to the pair R1-R2. Moreover, we applied a current injection to area R1 (Fig 2B) in order to test if CURBD is also able to detect not only information transfer - and therefore quantify functional connectivity - in baseline conditions, but also capture enhanced information transfer following a current injection mainly affecting strongly interconnected areas (in this case R1 and R2). We chose to apply current injection in R1 in order to contrast functional and anatomical connectivity, by assessing the elicited responses in R2. If we had opted for applying current in R2, the responses elicited in R1 might in fact have solely been due to the strong anatomical connectivity from R2 to R1. As expected, a current injection in R1 increased the overall firing rate in all regions (since all regions are interconnected), but significantly more strongly in R2, in line with the stronger interconnection between R1 and R2. (Fig 2B). When fitted to simulated data taken from 20 neurons per area, CURBD learned to reproduce the simulated data, for what pertains to both single unit activity (Fig 2C-left) and population-level activity patterns (Fig 2C-right).

We then proceeded to test the reliability of CURBD in estimating inter-areal connectivity, when considering a minimal number of neurons (one per area, for a total of 4 neurons). Here, we chose one neuron per area and applied the CURBD algorithm 1000 times (with a different random seed in each iteration), in order to determine if the connectivity estimates were consistent across different runs of the algorithm. We first applied this approach to the baseline activity (500 ms of spontaneous activity, Fig 3A), obtaining - as expected - significantly higher levels of connectivity estimates between R1 and R2 compared to all other pairs of regions (p<0.001, one way ANOVA with post-hoc Tukey correction). Therefore, CURBD applied to spontaneous network activity reflected the strength of anatomical connectivity between areas.

**Figure 3.**
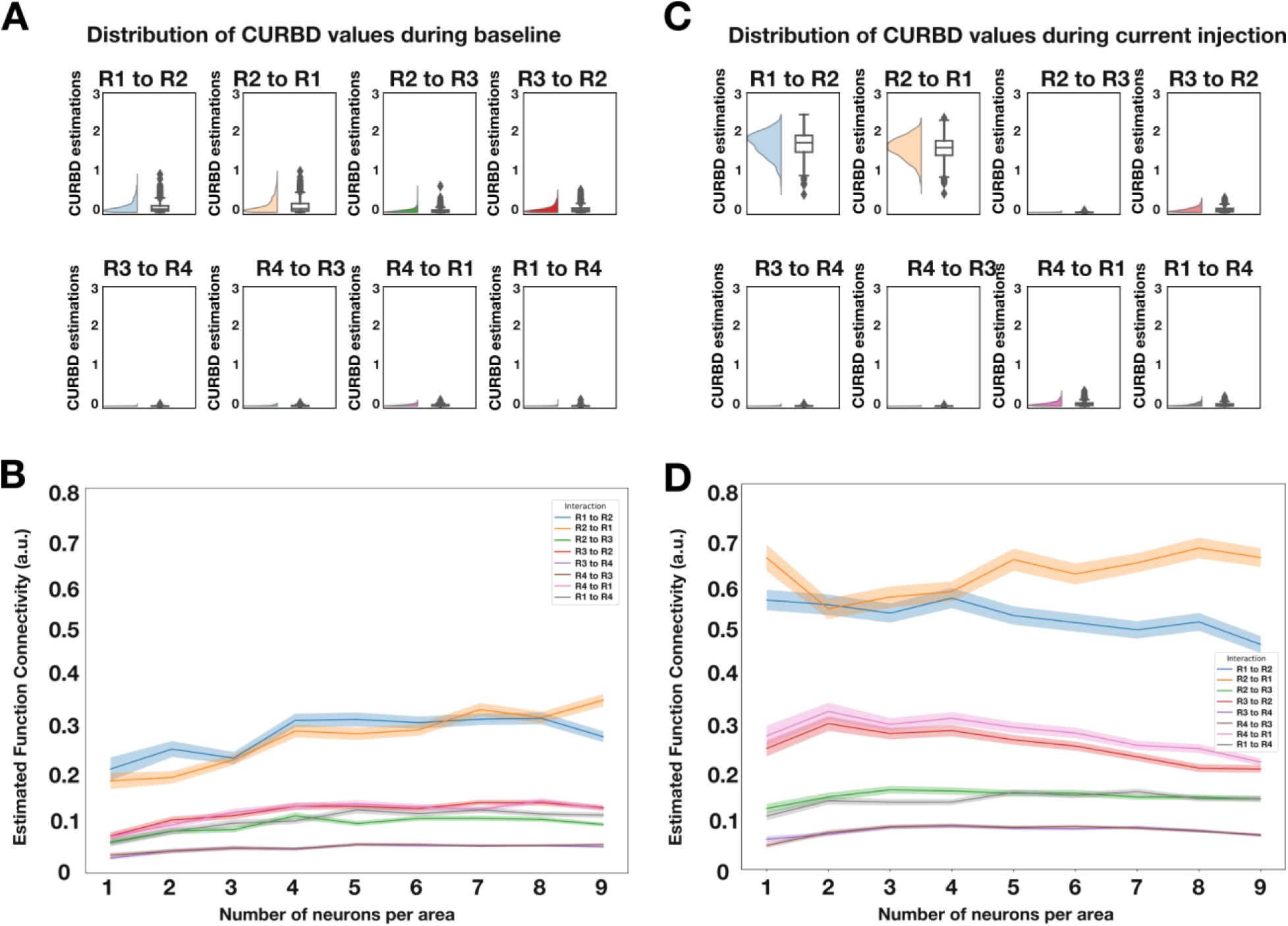
CURBD-based estimates of connectivity reflect both anatomical as well as effective connectivity and are valid already with a very limited sample size. **A.** For a selection of one unit per area (total of four units), we estimated the functional connectivity between the four different areas generated by the neural spiking model (Fig 2) across 1000 different runs of CURBD. For each iteration, we applied the CURBD algorithm to the same selection of units but using a different random seed, in order to observe if CURBD would be prone to spurious results when estimating connectivity with a low number of units. Estimated values during baseline (i.e spontaneous activity) are significantly higher between the pair R1-R2, matching the anatomical connectivity of the model. **B.** Varying the number of units per region leads to similar connectivity values. In order to assess if the results shown in panel A are dependent on the specific four units we chose for the initial simulation, we ran another set of tests. First, we varied the number of pooled cells from the model per region. Then, for each N selected units per region, we proceeded to randomly select N different cells on each run. We ran this test 500 times for each number of N units. Each line represents the average connectivity values observed after all the simulations, with SEM as the shaded area. **C**. Same as A, but for a period of activity after current injection. Note how the level of CURBD-estimated connectivity is significantly higher between R1 and R2 (and also higher compared to values observed during spontaneous activity), in line with the observed patterns of activity. **D.** Same as B, but for a period after current injection.

Moreover, we also tested if this result was specifically dependent on the 4 neurons we chose to apply the algorithm in the previous test, as well as on the number of neurons included in the computations. Specifically, we varied the number of neurons provided to CURBD per region (e.g 1 neuron per area, 2 neurons per area). For each selection of the number of neurons we ran the CURBD algorithm 500 times, each time randomly selecting a set of neurons from each of the simulated regions. The estimated level of connectivity remained rather constant independently of the number of neurons (Fig 3B), showing that, even when including very few neurons, the strength of functional connectivity between areas can be effectively estimated via CURBD (see also (Perich et al., 2020)).

Then, we applied the same approach to simulated neural activity after current injection in R1. This enabled us to test to what extent CURBD captures not only anatomical connectivity (that will vary across brain states except for some minor effects due to ongoing synaptic plasticity), but also effective connectivity, i.e. information transfer between regions. Indeed, during a 500 ms period of current-induced enhanced activity, CURBD estimated a level of functional connectivity that is generally higher in magnitude (Fig 3C) in comparison to baseline, and in particular between R1 and R2, in line with the observed changes in spiking activity. Specifically, connectivity between R1 and R2 was significantly stronger than that between all other pairs of regions (p<0.001, one way ANOVA with post-hoc Tukey correction). Moreover, when varying the number of provided units per region, analogously to what we observed for spontaneous activity, there were no strong variations on the estimated levels of connectivity (Fig 3D).

Overall, these results indicate that CURBD can reliably capture information transfer between brain regions even when few neurons can be recorded. Of relevance, the inclusion criterion we set for the in vivo recordings (see the Sample Size section) far exceeds the minimum number of one neuron per region observed in simulations to take into account physiological variability in neural activity (see also the results of the pilot analysis presented in Fig 5). Besides validating CURBD, this analysis clarifies that this method estimates effective connectivity between areas, based on how activity in one brain region determines patterns of spiking activity in another region. This is of course influenced by the underlying anatomical connectivity, but is also dependent on the actual engagement of individual brain regions in information processing and communication. However, as the term effective connectivity is often implied to include the estimation of causal effects - that we cannot estimate in the absence of an interventional approach using e.g. optogenetics - we instead stuck to the term functional connectivity in the rest of the manuscript.

### Statistical plan

#### Hypothesis testing

For all the defined hypotheses, the dependent variable is, as discussed above, the estimated functional connectivity between pairs of neurons located in different areas (see Estimation of connectivity for more details). Specifically, for each 5 s epoch, we computed the current estimation over the whole epoch for all pairs of neurons located in different areas (Fig 4A). This gave us an estimate of the inter-areal connectivity between all the recorded areas in a given 5 s epoch (Fig 4B). This represents the sample unit for all the statistical analyses described in the following sections. We tested the following hypothesis regarding the functional connectivity among brain regions in NREM and REM sleep, respectively, independently:

**Figure 4.**
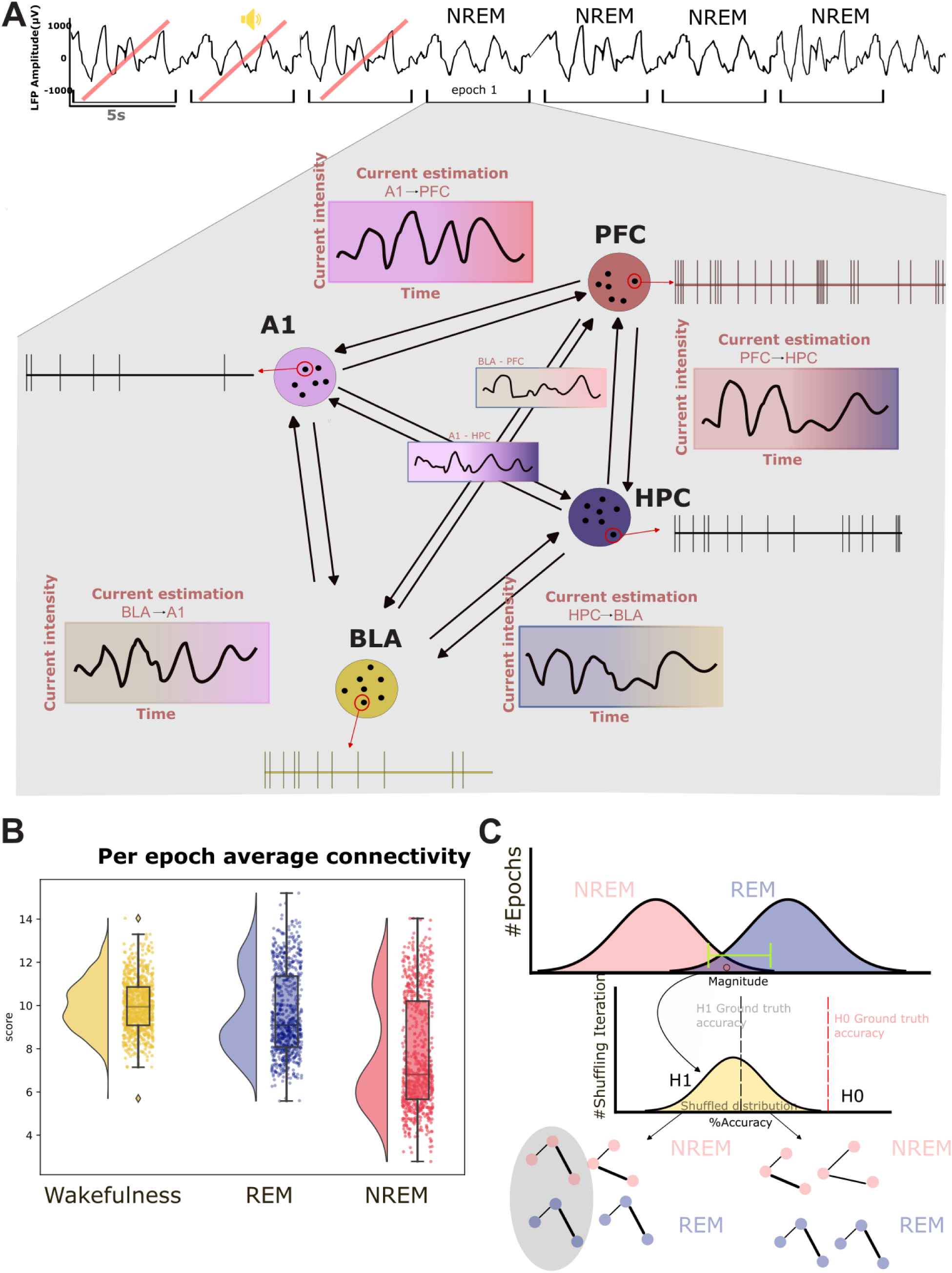
Estimation of connectivity and hypothesis testing. **A**. For each valid sleep epoch, single unit activity of all pairs of inter-region responsive neurons will be provided to the CURBD network. For each of these pairs we will calculate a 5s time series of current intensity, which is CURBDs’ measure of connectivity. **B.** By averaging, for each epoch, each of the current estimation time series, we will obtain a single score representing the measured connectivity on that epoch, between neurons of different regions.. **C.** We hypothesize that, despite global differences, a significant number of NREM epochs might contain connectivity values comparable to REM. For each epoch, through a leave-one-out approach performed on the higher tail of the NREM pairwise connectivity distribution, a classifier is trained to distinguish between NREM and REM epochs’ connectivity. The same procedure will be done by comparing NREM and wakefulness. By observing the distribution of the classifier performance, we can confirm H1 if the classifier returns a low ground truth accuracy. Similarly, if it returns a high accuracy, we can confirm H0. Importantly, this procedure will also take in account the shape of the network, instead of focusing solely on the average global values of all pairs.

#### Hypothesis 1 (H1)

Hypothesis 1 (H1): A significant fraction of NREM epochs displays a structure of interareal FC that is indistinguishable from what is observed on average in REM and wakefulness. Null hypothesis for NREM (NREM_H0): less than 5% of NREM epochs exhibit a level of FC comparable to that observed in REM and/or wakefulness; for those NREM epochs with average FC values comparable to those observed in REM and wakefulness, a structure that is different from that observed in REM and/or wakefulness is observed. Of relevance, we separately compared NREM to wakefulness and REM, but have no a priori hypothesis about any possible differences between wakefulness and REM.

In view of previous studies (Olcese et al., 2016; Olcese, Bos, et al., 2018), we expect that FC will not significantly vary for pairs of neurons located in the same brain region. Therefore, we focused on pairs of neurons located in different regions. We analyzed the data obtained in different experimental conditions (Habituation, Fear conditioning and Fear test). We tested H1 at the level of individual recording sessions and we expected the effect to be present independently from experimental conditions taking place in the awake state. To test H1, we first averaged all FC values obtained in a given epoch for all inter-areal pairs of neurons. This allowed us to compute a global value of inter-areal FC at single-epoch resolution. Based on previous studies (Massimini et al., 2005; Olcese et al., 2016; Olcese, Bos, et al., 2018), we expected that the global value of inter-areal FC is, on average, lower during NREM than in either REM and wakefulness. State-dependent functional connectivity between cortical areas is expected to drop during NREM sleep, in comparison to wakefulness, both in view of empirical evidence (Massimini et al., 2005; Storm et al., 2017) as well as theoretical considerations (Koch et al., 2016; Tononi et al., 2024). In contrast, there is evidence that the subcortical-cortical communication (e.g from the hippocampus to cortical areas), can be preserved or even enhanced during NREM sleep (Chen & Wilson, 2023; Ji & Wilson, 2007; Klinzing et al., 2019; Louie & Wilson, 2001). These specific state-dependent functional connectivity changes across cortical and subcortical regions have, however, been shown to be dependent on the connectivity measure and time scale under consideration (Olcese et al., 2016; Olcese, Bos, et al., 2018). In view of this tight relationship between the specific details of the method that is used and the results that are obtained, we focused on quantifying how CURBD-estimated connectivity varies across brain states, in terms of average values and structure, without considering whether distinct effects might occur based on which specific pairs of areas are considered. Overall, based on previous studies (Massimini et al., 2005; Olcese et al., 2016), we expected a global drop of connection strength during NREM sleep in comparison to REM and wakefulness. Nevertheless, we cannot exclude that connection strength between some areas (e.g. between cortical areas and hippocampus) might follow different trends. This, however, would not be a problem for what pertains testing our hypotheses, since we investigated if the FC structure varies within brain states, and not specifically how it varies.

We tested if FC strength varies across brain states using a repeated measurement ANOVA with post-hoc Tukey correction (alpha: 0.05), or a corresponding non-parametric test in case the assumptions of normality are not satisfied. This is a verification that the FC values we obtained conform to the expected global changes across brain states, and is a prerequisite for testing H1. In case this analysis does not confirm the expected results, we will not pursue H1 and only present the results for average FC values.

The next step was to determine, within individual recording sessions, the fraction of NREM epochs with a global level of inter-areal FC high enough to be comparable with the values observed in REM and wakefulness (H1, Fig 4C). To this aim, we first computed the fraction of NREM epochs with an average inter-areal FC (aiFC) in the top 25th percentile for NREM and in any case higher than the 5th percentile of the distribution of aiFC values in REM (wakefulness). In view of the pilot results we have obtained (see the “Pilot Results”) section, this allowed to select NREM epochs whose aiFC values, irrespective of the source of the variability in FC values, are likely to be in the range observed during REM (or wakefulness) rather than NREM. If, nevertheless, the fraction of NREM epochs with aiFC values higher than the 5th percentile of the distribution of aiFC values in REM (wakefulness) is lower than 5% in more than 50% of the recording sessions, we would conclude that NREM_H0 cannot be rejected. Otherwise, we would test, in the recording sessions showing a significant overlap, if the NREM epochs with a high aiFC simply reflect a NREM network architecture with stronger FC values, or rather include epochs with a REM-like (wakefulness-like) network architecture (H1).

To test H1, we trained a classifier (support vector machine) to discriminate, on the basis of the values of individual directed connectivity values between inter-areal pairs of neurons, whether NREM epochs in the high tail of the aiFC distribution - high-aiFC, defined as described in the previous paragraph - are classified as either NREM or REM (wakefulness). The classifier was trained on high-aiFC NREM epochs and on all REM (wakefulness) epochs using a modified leave-one-out cross-validation scheme in which only the NREM epochs in the high tail of the aiFC distribution are individually tested. This procedure delivered a ground truth accuracy value for the classification of NREM vs. REM (wakefulness) epochs with similar values of aiFC. To test if this accuracy value reflects H1 or NREM_H0, we repeated the cross-validation procedure by shuffling the labels of NREM and REM (wakefulness) epochs with comparable values (i.e., separately within 1 out of 10 equipopulated bins in which the range of REM (wakefulness) aiFC values will be subdivided). More precisely, assuming there are N NREM epochs with aiFC values in the high tail of the distribution, we repeated the leave-one-out scheme by identifying N REM (wakefulness) epochs with values within the range of aiFC values observed in the selected NREM epochs. At each iteration of the cross-validation, 1 NREM epoch was left out for testing. For the other N-1 epochs, sleep stage labels were randomly shuffled with those of the selected REM (wakefulness) epochs with corresponding aiFC range (separately for each of 10 equipopulated bins in which the range of REM (wakefulness) aiFC value will be subdivided, i.e. each bin had a width of ten percentiles of the distribution of aiFC values). The shuffling procedure was repeated 1000 times. This shuffling procedure allowed us to obtain, for each recording session, 1000 estimates of classification performance for NREM and REM (wakefulness) epochs not only with comparable aiFC values, but also indistinguishable network architecture. In case the ground truth accuracy value falls within the 95% confidence interval of the shuffled estimates, this meant that there was, within NREM epochs, a significant fraction of epochs whose inter-areal network architecture is indistinguishable from that observed in REM (wakefulness) - Fig 4C - therefore supporting H1. A p-value was calculated as the fraction of instances in which a classifier trained on the shuffled dataset shows an accuracy greater than ground-truth classification accuracy.

The procedure described above was repeated three times, respectively on NREM epochs with FC values in the top 25%, 15% and 5% of observed NREM values. This allowed us to test H1 as a function of the strength of aiFC observed in NREM. If ground truth accuracy value falls within the 95% confidence interval of the shuffled estimates for a given range of high-aiFC values (top 25%, 15% and 5%), this would be seen as a confirmation of H1 and rejection of NREM_H0. If the ground truth accuracy is instead higher than the 95% confidence interval of the shuffled estimates - separately estimated for each of the three ranges of high-aiFC (top 25%, 15% and 5%) - we would not be able to reject NREM_H0.

To rule out the possibility that failure to reject H1 might stem not from a confirmation of our hypothesis, but from other aspects, such as an insufficient sample size, we trained a classifier to discriminate NREM epochs in the 5th percentile for FC strength from REM (or wakefulness) epochs in the 95th percentile for FC strength. As these epochs are those assumed to differ the most in terms of FC strength, we expected that a classifier would not be able to correctly (i.e. above chance) discriminate to which brain state each epoch belongs only if sample size is too low or if other aspects (e.g. noise in CURBD estimates) are insufficient to obtain sufficiently high accuracy. Ground-truth classification accuracy was compared to accuracy of a classifier trained after randomly shuffling the labels of NREM and REM (or wakefulness) epochs. If no difference was observed between the two (i.e. if ground-truth accuracy falls within the 95% confidence interval of accuracy for classifiers trained on the shuffled datasets) we would exclude the session.

Finally, we combined the results - separately for each of the three high-aiFC ranges under consideration - across multiple recording sessions. In case the sample size analysis shows that we have enough power to draw meaningful conclusions from different types of sessions (e.g. fear conditioning vs. habituation), we performed this analysis. Otherwise, we would only compute a combined p-value for all sessions (via Fisher method). If the combined p-value across all sessions is larger than 0.05, we interpreted this as a confirmation of H1 and rejection of NREM_H0 (see Fig 4B). Conversely, a combined p-value lower than 0.05 would not allow us to reject NREM_H0. In order to control for the possibility of individual animals strongly influencing the outcome of our analysis, we would also apply Fisher’s method to separately combine p-values within each animal. We reported the combined p-values per animal and verified that comparable results can be obtained across all animals. In view of the limited number of animals, in case we find that results differ between animals, we would conclude that our dataset is underpowered.

#### Hypothesis 2 (H2)

Hypothesis 2 (H2): A significant fraction of REM epochs displays a structure of interareal FC that is comparable to what is observed on average in NREM. Null hypothesis for REM (REM_H0): <5% of REM epochs generally exhibit a level of FC comparable to that observed in NREM; for those REM epochs with average FC values comparable to those observed in NREM, a structure that is different from that observed in NREM is observed. To test this hypothesis (analogous to H1, but for REM sleep), we followed the exact same procedures as for H1. We trained a classifier (support vector machine) to discriminate, on the basis of the values of individual directed connectivity values between interareal pairs of neurons, whether REM epochs in the low tail of the aiFC distribution - those epochs whose aiFC value is in the bottom 25th, 15th or 5th percentile of aiFC values for REM epochs (each of these three ranges will be separately analyses, similarly to H1) and in any case lower than the 95th percentile of aiFC values for NREM epochs - are classified as either NREM or REM.

#### Sampling plan

For this study, we utilized an existing dataset (see Table 5), in light of the ethical principle of reduction in animal research. This significantly lessens the number of animals used by avoiding unnecessary replication of data collection. Moreover, leveraging existing data promotes efficient use of resources and prevents potential biases introduced during data collection. In case our sample size analysis concludes that the proposed dataset does not have enough statistical power to draw the conclusions we would add different recording sessions that were obtained from the same experiment. Here we propose carrying the investigation on the first three days from a fear conditioning and extinction paradigm, but two additional recording sessions (extinction training and extinction test) are available in case it is needed. These sessions have not been pre-processed yet but may yield additional data, pending quality control.

**Table 5:** Dataset description.

| Animal ID | Session type | BLA (number of units) | PFC (number of units) | A1 (number of units) | HPC (number of units) | NREM (number of 5s epochs) | REM (number of 5s epochs) | WAKE (number of 5s epochs) | Enough epochs |
| --- | --- | --- | --- | --- | --- | --- | --- | --- | --- |
| r14 | Habituation | 7 | 7 | 16 | 1 | 1658 | 541 | 665 | Yes |
| r14 | Fear Conditioning | 6 | 9 | 14 | 3 | 1475 | 637 | 853 | Yes |
| r14 | Fear Test | 8 | 11 | 11 | 5 | 1261 | 661 | 1464 | Yes |
| r16 | Fear Conditioning | 6 | 6 | 4 | 5 | 2641 | 804 | 1169 | No |
| r20 | Habituation | 7 | 11 | 8 | 4 | 2556 | 772 | 1035 | Yes |
| r20 | Fear Conditioning | 4 | 9 | 10 | 4 | 2511 | 1290 | 2129 | Yes |
| r20 | Fear Test | 8 | 5 | 1 | 5 | 2065 | 752 | 1382 | Yes |

Our dataset was collected through multi-area tetrode recordings (Olcese et al., 2016; Olcese, Bos, et al., 2018). This is a technique that enables the simultaneous recording of a more limited number of neurons, when compared to recently introduced techniques (e.g. chronic Neuropixels probe recordings (van Daal et al., 2021)). Furthermore, we are only able to determine in which brain regions recorded neurons were located, but not in which area subdivision (e.g. cortical layer), nor to which neuronal subpopulation a neuron belongs (e.g. which type of interneuron).

In view of the relatively low number of recorded neurons, we have decided not to subdivide them in subpopulations - e.g. between putative excitatory or inhibitory neurons based on the action potential waveforms (Olcese et al., 2013; Oude Lohuis et al., 2022). Since our analysis is based on comparing patterns of inter-areal FC (i.e., averaged across pairs of neurons), we included both putative single units and high-quality MUA (see also the “Spike Detection” section). Furthermore, as explained earlier, we did not aim to quantify or interpret connectivity patterns between individual neurons. Instead, we only quantified the average connection strength measured for neurons located between brain regions. This approach, that we successfully employed in previous studies (Olcese et al., 2016; Olcese, Bos, et al., 2018), would thus prevent the risk of drawing unwarranted conclusions from a limited number of recorded neurons. Therefore, this study only allowed us to draw conclusions about how brain states modulate pairwise functional connectivity between areas. A microcircuit-level characterization of such modulation may be performed at a later stage, based on the results that we obtained. Furthermore, all the four areas from which recordings were done were included. In fact even if not all the recorded areas are expected to contribute to establishing the level consciousness, previous studies showed that inter-areal connectivity is modified across most areas (Olcese et al., 2016; Olcese, Bos, et al., 2018). Thus, independently of whether connectivity between areas is depressed (as is expected between cortical areas during NREM sleep compared to wakefulness) or enhanced (as may occur between hippocampus and cortex during NREM compared to wakefulness), we would nonetheless be able to use the recorded data to address our key hypothesis about the heterogeneous nature of brain states. Crucially, as we discussed in the section “Sample size”, the number of neurons recorded in each session is estimated to be sufficient to obtain valid CURBD estimates. Importantly, in view of the fact that different set of neurons were recorded during each recording sessions (as tetrodes were independently advanced through brain tissue by at least 250 μm), but also following the typical convention in the field (Bos et al., 2019; Dorman et al., 2023; Olcese et al., 2016; Olcese, Bos, et al., 2018) and standard neuroscientific practice, each recording session should in fact be considered a separate observation (Asaad & Sheth, 2024). Thus, even if data was collected from only three animals, we have in fact access to at least seven separate recording sessions, that we may extend based on the quality of extra recording sessions collected during fear extinction sessions if necessary (see also earlier sections).

#### Exclusion Criteria

We applied two key exclusion criteria in our analysis. First, following our pilot analysis on the effectiveness of CURBD in fitting the collected data as a function of the number of neurons (see the Sample Size section), we excluded every recording session where the total number of neurons is less than 10. Second, we excluded epochs in which less than two neurons were firing, due to their potential to generate noisy and statistically insufficient data. Third, we excluded epochs where sound stimulation occurred, along with the immediately adjacent epochs, to minimize potential confounding effects and to focus exclusively on spontaneous activity changes during sleep. Fourth, in cases in which there are no significant values for interareal connectivity across all considered brain states, we interpreted this as an indication of no observable communication between the considered areas. We excluded the corresponding session from further analysis.

#### Sample size

In view of the nature of the analyses we performed, we are not able to make an estimation of the effect size we obtained. For this reason, we could not perform a classical power analysis, but instead focused on determining if we have a sufficient sample size (number of neurons, epochs, sessions) to draw significant conclusions. We have separately addressed this issue for what pertains a) the minimum number of neurons required by CURBD to obtain reliable estimates of FC, b) the number of epochs required to reliably estimate different FC structure between brain states and c) the number of recording sessions needed overall to obtain reliable results.

### Number of neurons in each recording session

Even if CURBD does not theoretically require a minimum amount of neurons to provide meaningful current estimations (Perich et al., 2020), we nonetheless performed a pilot analysis meant to determine which recording sessions contain a number of neurons high enough to obtain a reliable CURBD-based estimate of inter-areal functional connectivity.

In order to avoid performing any analysis on the portion of the dataset that we aim to use in the following phase of the project, we decided to focus on epochs that we planned to exclude from the main analysis. In detail, our dataset included recordings performed in rats that were also exposed to auditory stimuli during sleep. In the main analyses we focused on spontaneous spiking activity, and removed all epochs that contain sound exposure. For this additional analysis we instead extracted all the epochs during one single example REM sleep session in which a specific sound (CS+) was played (Fear test, table 5). In other words, we only used epochs that were excluded from the main analysis, from one recording session only. Then, we subsampled data from the selected recording session by randomly selecting a number of neurons ranging from 4 to 24. For each value of the number of neurons, randomly sampled 500 different combinations of neurons, and for each of these combinations we obtained CURBD estimates. Then, we quantified the partial variance (pVar) that each fitted model was able to explain about the recorded data. This analysis revealed that, when at least 10 neurons were included in a subsampled dataset, any additional neuron did not improve the variance explained by the model (see Fig 5). Therefore, we excluded sessions that did not have a minimum of 10 units, as they would lead to suboptimal CURBD estimates.

**Figure 5.**
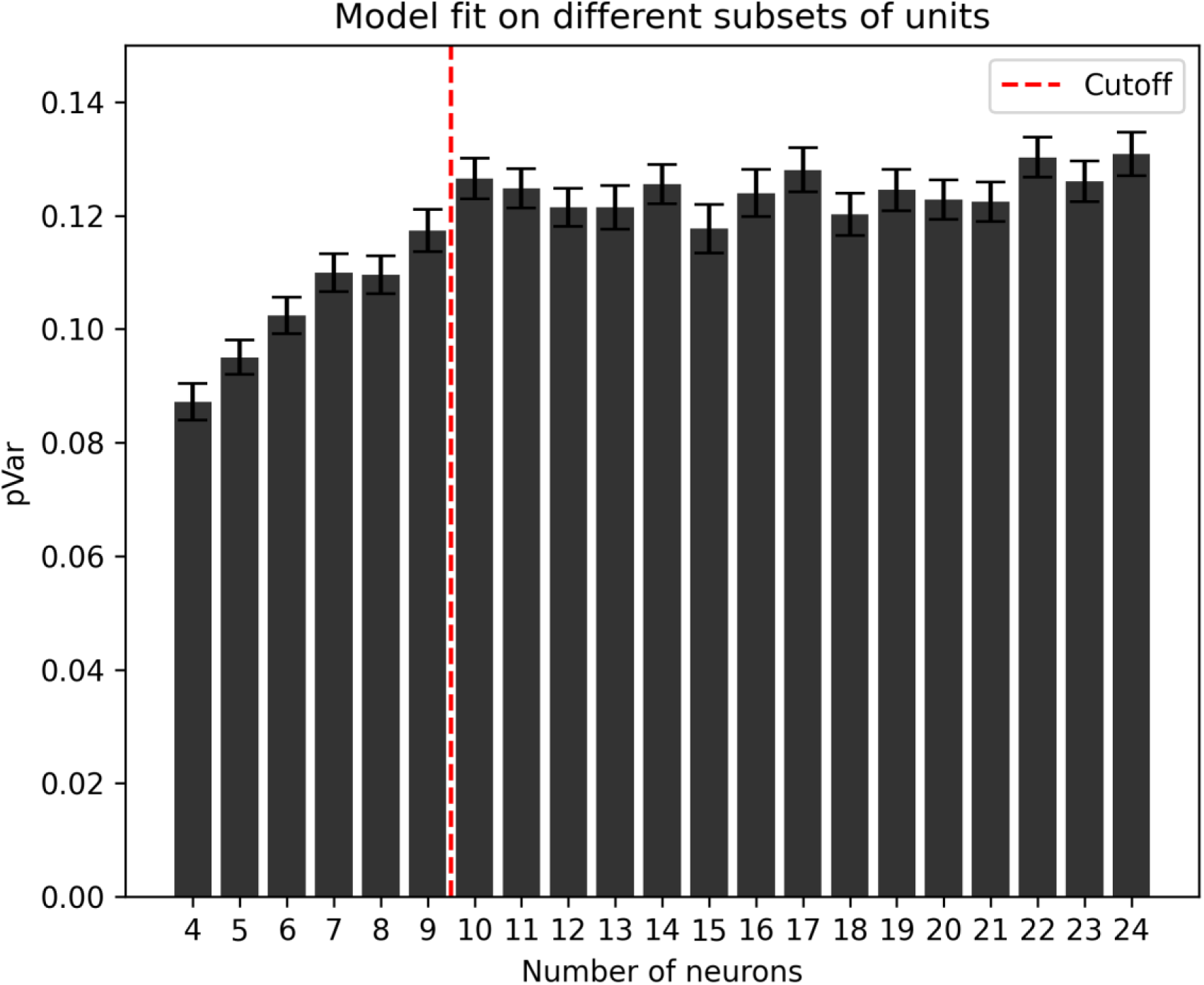
Model goodness of fit as a function of the number of recorded neurons. Systematic testing of partial variance (pVar) explained by CURBD models fitted on a subset of neurons from a sample dataset (500 resamplings per number of neurons) shows how pVar varies as a function of increasing number of neurons. We analyzed which bins were significantly different from the adjacent ones (where a bin indicates the number N of neurons included in the model). This analysis revealed that only two adjacent bins were statistically different from each other: the bins N=9 and N=10 (t-test, p = 0.018) and the bins N=13 and N=14 (t-test, p = 0.037). Moreover, to further explore whether the goodness of fit of the model showed a plateau after a certain number of neurons is included, we took the subset of neurons with the highest explained variance pVar (N=24) and compared all the other bins to it. This analysis showed us that all statistical comparisons from the subset N= 10 until N = 23 were not significantly different (Bonferroni corrected t-test, alpha = 0.01). All bins from N = 4 until N = 9 were significantly different from the N=24 bin (Bonferroni corrected t-test, p <0.001). All recording sessions with less than 10 neurons (red dashed line: cutoff) will therefore be excluded from further analyses.

### Number of epochs in each recording session

For what instead pertains to the number of sleep epochs in each session, the availability of enough data to potentially reject the null hypothesis is ensured by the shuffling procedure we discussed earlier. Specifically, our assessment is focused on the availability of a sufficiently large dataset to train the classifier used in testing H1/2. To test this, we harnessed the classifier’s ability to discriminate between two brain states by using a similar leave-one-out scheme as that described in the “Hypothesis testing” section. The classifier was trained on all the available epochs for two states (e.g. NREM and REM) and the performance was tested via a cross-validated leave-one-out scheme using all epochs (and not just the epochs in the tail of the aiFC distribution. The performance of the classifier was tested against that obtained after shuffling the labels of behavioral state (same procedure as described in the “Hypothesis testing section”. If the classifier is able to discriminate which epoch belongs to which state with an accuracy significantly higher than chance (i.e. larger than the 95th percentile of accuracy values obtained on the shuffled training sets) we concluded that the number of epochs is sufficient to apply the proposed approach. This verification was done for all recordings and between all pairs of brain states. Any session and pair of brain states with a classification accuracy not higher than chance was excluded from further analyses. The results of this analysis was reflected in the stage 2 manuscript by adding a column to table 5 with the indication, for each recording session, of whether a sufficient number of epochs to compare different brain states is present or not.

Thus, by comparing the ground truth classification accuracy to the value obtained with the shuffling procedure we tested if we can reject or not H0. In case H0 can be rejected, this will necessarily mean that we have enough epochs to compute valid classification estimates and that H1 (or H2) are verified. Otherwise, we would not be able to make any claim about H1 or H2.

### Number of recording sessions

Finally, we developed a simulation-based approach to estimate the power of the meta-analysis performed when combining p-values obtained from individual recording sessions (Fig 4B). This is necessary because, in view of the novel nature of our research question and methodology, we are unable to make any a priori assumption of the effect size and variability of the effect we would see. Therefore, to avoid any risk of circularity (Hoenig & Heisey, 2001), a simulation-based power analysis was performed based on a set of pilot analyses run on 3 recording sessions (Fridley et al., 2010). First, for each of these sessions, we identified the classification accuracy corresponding to a p-value of 0.05. Then, we randomly resampled data (with replacement) from these 3 sessions to generate surrogate datasets consisting of 7 sessions (same number of sessions as in our dataset). For each session in a surrogate dataset, we computed the p-value associated to the accuracy level corresponding, in the original recording session from which the data was drawn, to a p-value of 0.05. For each surrogate dataset a combined p-value was computed via Fisher’s method. The procedure was performed 1000 times to compute a statistical power. If power results to be higher than 80% we continued with the analysis, otherwise we concluded that we do not have sufficient data to draw any conclusion, i.e. that our dataset is underpowered. In case the dataset is deemed as underpowered, we can expand it with other remaining sessions of our behavioral paradigm (see also the Sampling plan). Conversely, this analysis might show that we have enough data to quantify possible differences between the various types of sessions (e.g. fear conditioning independently from baseline controls). In this case we analyzed the different types of sessions separately.

#### Limitations

The proposed study aims to re-use an existing, previously collected dataset. This prevents the need to perform novel experiments and is thus in line with the well-recognized need to reduce the number of animal experiments. Nevertheless, this approach also has some downsides. First of all, we were not able to collect novel data. Therefore, a risk is that the available sample size might result to be insufficient to draw statistically significant conclusions. In particular, we are aware that the data was collected from only three animals, each undergoing a set of experimental conditions unrelated to the scientific objective of the current study. This sample size is comparable to what is often done for studies involving multi-area tetrode recordings (see also the “Sampling plan” section for more details). However, we cannot exclude that distinct animals or experimental conditions might show different results. Furthermore, we are also aware that our recordings yielded a low number of neurons, especially when compared to novel technologies such as Neuropixels probes. However, these methods were not yet available when the experiment we analyzed here was performed. Based on the modeling study we performed, we are confident that the CURBD method yielded reliable results even with a limited number of neurons. However, we expect that our study will be followed up by new investigations involving large-scale recordings that will be able to further characterize how FC is modulated by brain states.

### Pilot Analyses

In order to perform a preliminary test of the analytical procedures described here, we have performed a pilot analysis on single units recorded from one randomly selected recording session (animal r20, fear test session). First, we aimed to test if the assumptions underlying the ability to perform H1 (i.e. of higher inter-areal FC values - in absolute value - in wakefulness compared to NREM sleep) were verified. This was verified (Fig 6A; p<0.05, Mann-Whitney U test), although we also observed a considerable overlap between FC values observed in wakefulness and NREM. Such a high overlap in FC values could reflect a limited change in FC values between wakefulness and NREM, but also be indicative of a low reliability of CURBD-estimated FC. To better understand this, we first computed inter-areal FC strength while excluding connections to and from the hippocampus - since cortico-hippocampal communication is known to be preserved in NREM sleep compared to wakefulness (Olcese et al., 2016; Olcese, Bos, et al., 2018); we observed that the overlap between FC values - quantified by looking at the fraction of NREM FC values falling within each quartile of the distribution of wake FC values - was lower compared to when all the recorded areas were included (Fig 6B). We then focused on epochs in which the highest difference in FC between wakefulness and NREM could be expected. Specifically, we focused on the 10% of NREM epochs with the highest slow wave activity, obtained by calculating the LFP power between 0.5 and 4Hz, and the 10% of wakefulness epochs with the highest motor activity, obtained by calculating the area under the curve of accelerometer activity in each epoch. As expected, the distributions of FC values were more different (both in terms of average value and overlap between the distributions) compared to when we included all epochs in the analysis (Fig 6C), an effect that was even more prominent when we excluded connections to and from the hippocampus - Fig 6D. Conversely, and also in line with previous studies (Olcese et al., 2016; Olcese, Bos, et al., 2018), no significant difference was observed between wakefulness and sleep for intra-real FC strength (Fig 6E). Finally, and also in line with our assumption, a significant difference was observed for inter-areal FC coupling between NREM and REM (with stronger FC during REM - Fig 6F; p<0.05, Mann-Whitney U test).

**Figure 6.**
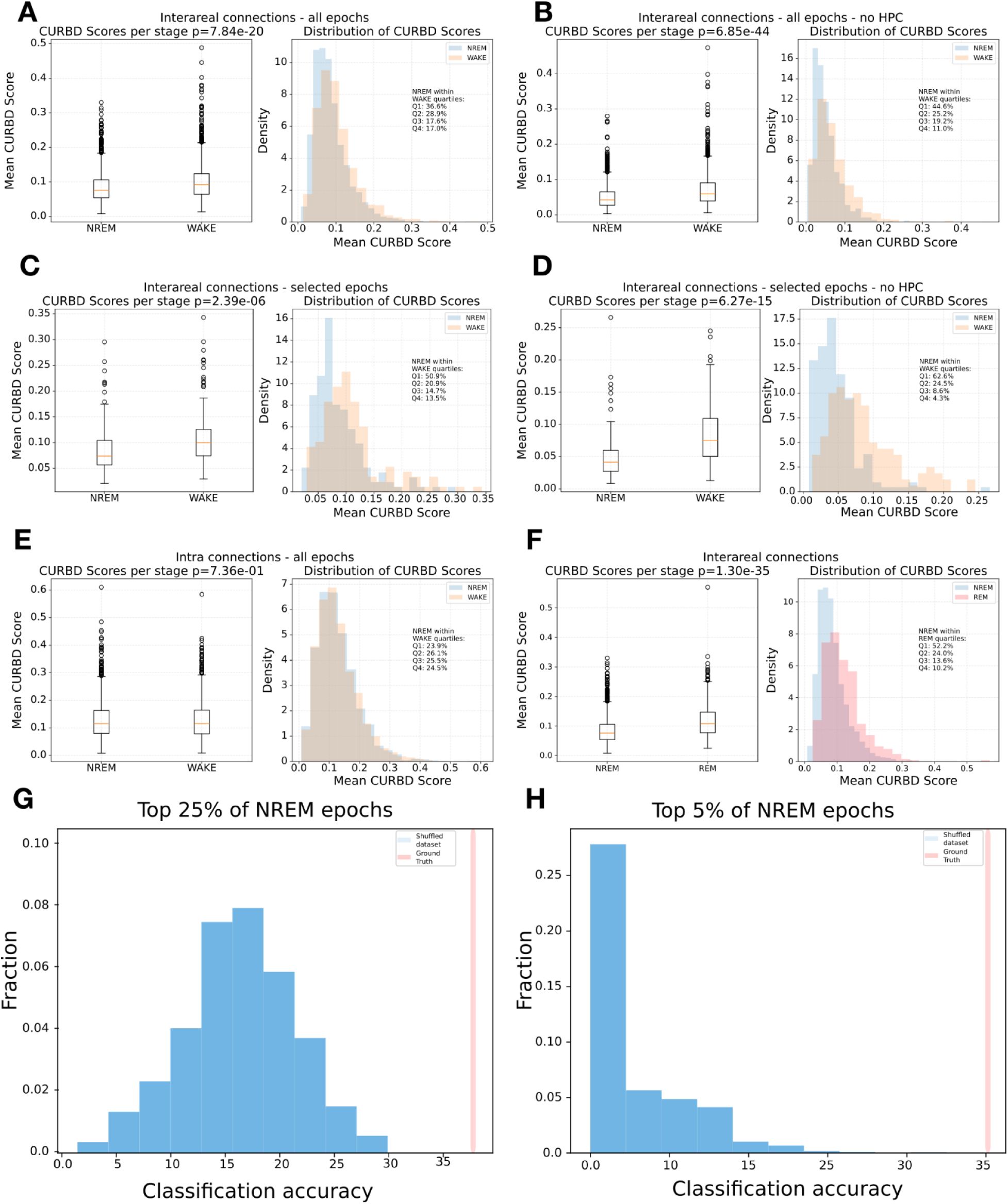
Pilot results for a randomly selected recording session. **A.** Left: Box plots showing the distribution of average inter-areal (aiFC) values for NREM vs. wakefulness epochs. Right: Histograms of the distribution of aiFC values for NREM vs wakefulness, with indicated the fraction of NREM epochs falling within each quartile of the distribution of aiFC values in wakefulness. **B.** Same as *A*, but excluding connections to and from the hippocampus. **C.** Same as *A,* but only for selected NREM epochs with the top 10% of SWA values and wakefulness epochs with the top 10% of detected motor activity. **D.** Same as *C*, but excluding connections to and from the hippocampus. **E.** Same as *A*, but for intra-areal connections. **F.** Same as *A*, but for NREM vs. REM. **G.** Classification accuracy for a classifier trained to discriminate high-aiFC NREM epochs from wakefulness epochs. Red: Ground-truth accuracy. Blue: Histogram of the accuracy obtained for classifiers trained on shuffled datasets. **H.** Same as *G*, but when considering NREM epochs with the top 5% of aiFC values.

Finally, in order to test the full analytical procedure to compare FC structure between brain states, applied the analytical procedure, as described in the “Hypothesis Testing” section to compare the FC structure between high-aiFC NREM epochs and wakefulness epochs, separately for the NREM epochs with FC values in the top 25% and top 5% (Fig 6G and 6H, respectively). In both cases, the ground truth accuracy is higher than the 95% confidence interval of accuracy values for classifiers trained on the shuffled dataset. This pilot analysis is important to gauge the range of accuracy values obtained for both ground truth and shuffled datasets (note the unbalanced nature of the dataset, with more wakefulness epochs compared to high-aiFC epochs). Although this pilot analysis reject H1 for this specific recording session (cf. Fig 4C), we refrain from drawing any conclusion.

## Results

### Exclusion Criteria

#### Interareal connectivity across brain states

The first step in our analysis pipeline assessed the effectiveness of the CURBD algorithm in identifying interareal connectivity across brain states. Experimental sessions showing no activity were to be excluded from further analysis, as this would indicate absence of communication between the target brain areas. However, all experimental sessions displayed functional connectivity (FC) values higher than 0, thus no sessions were excluded based on this criterion.

Anticipating one control step, we also tested the hypothesis that average FC should differ in magnitude across brain states. Six of the seven considered sessions showed statistically significant differences between sleep stages (awake, NREM, REM; p < 0.05, Kruskal-Wallis test; Table 6).

**Table 6:**
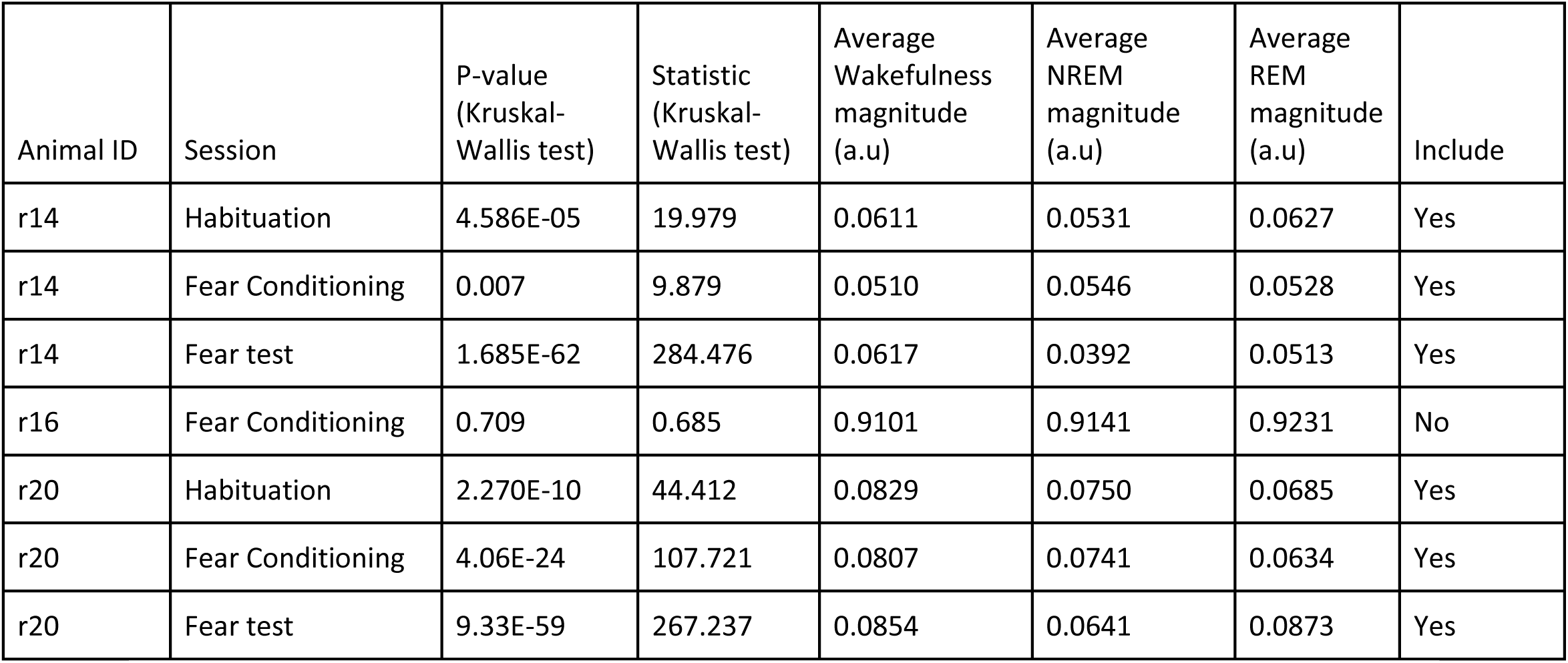
Statistical comparison of interareal functional connectivity magnitude across brain states (wakefulness, NREM, and REM) during different experimental sessions. The table shows the Kruskal-Wallis test statistics and corresponding p-values, along with average connectivity magnitudes (in arbitrary units, a.u.) for each brain state. Sessions were excluded in case the p-value was lower than 0.05.

| Animal ID | Session | P-value<br>(Kruskal-Wallis test) | Statistic<br>(Kruskal-Wallis test) | Average Wakefulness magnitude (a.u) | Average NREM magnitude (a.u) | Average REM magnitude (a.u) | Include |
| --- | --- | --- | --- | --- | --- | --- | --- |
| r14 | Habituation | 4.586E-05 | 19.979 | 0.0611 | 0.0531 | 0.0627 | Yes |
| r14 | Fear Conditioning | 0.007 | 9.879 | 0.0510 | 0.0546 | 0.0528 | Yes |
| r14 | Fear test | 1.685E-62 | 284.476 | 0.0617 | 0.0392 | 0.0513 | Yes |
| r16 | Fear Conditioning | 0.709 | 0.685 | 0.9101 | 0.9141 | 0.9231 | No |
| r20 | Habituation | 2.270E-10 | 44.412 | 0.0829 | 0.0750 | 0.0685 | Yes |
| r20 | Fear Conditioning | 4.06E-24 | 107.721 | 0.0807 | 0.0741 | 0.0634 | Yes |
| r20 | Fear test | 9.33E-59 | 267.237 | 0.0854 | 0.0641 | 0.0873 | Yes |

As expected, we observed a global drop of functional connectivity in NREM compared to wakefulness and REM (Fig 7A). Only one session (r16 fear conditioning) was excluded due to a lack of difference in FC magnitude across brain states. Following the statistical plan, we refrained from comparing FC amplitude values between specific brain states at this stage. However, we noted that not all sessions aligned with the expectation of a lower FC amplitude in NREM compared to wakefulness and REM (Fig 7B). This will be addressed in the next steps of the analysis pipeline.

**Figure 7.**
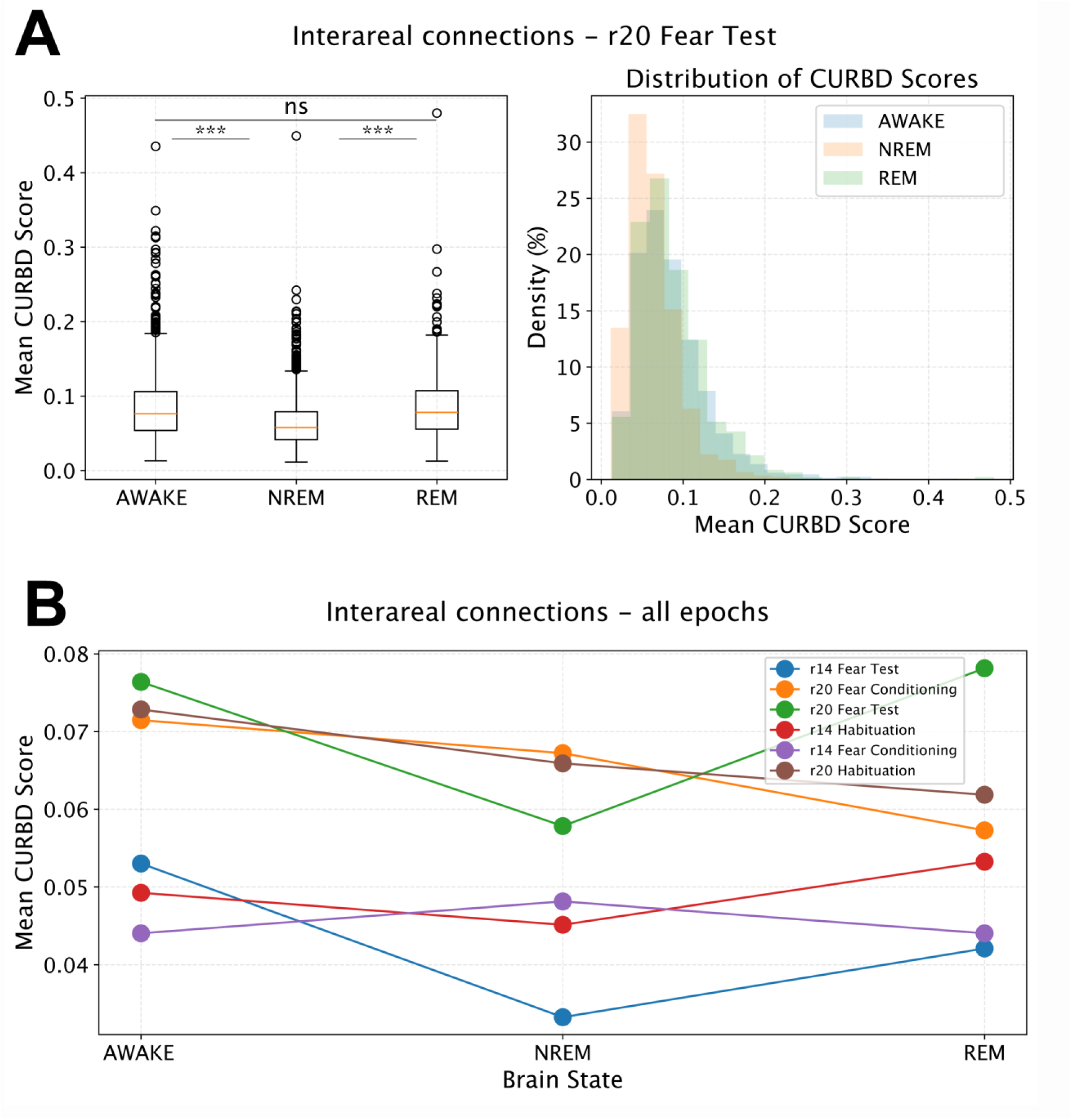
Interareal connectivity across brain states. **A.** Left: Box plots illustrate the distribution of mean functional connectivity (FC) magnitude values across all sessions, per brain state. Asterisks denote a statistically significant difference (p < 0.001, Kruskal-Wallis test) for the indicated tested pairs. ns: not significant. Right: Histogram displaying the overall distribution of FC magnitude values across all sessions. **B.** Mean FC values are presented for each session and each distinct brain state. Individual lines represent separate experimental sessions.

#### Number of epochs

Next, we assessed if the number of sleep epochs recorded per animal was sufficient to draw statistically meaningful conclusions. Here, separately per session and per relevant pair of brain states (wakefulness vs NREM, REM vs NREM; Fig 8A-B), we trained a classifier (support vector machine, with leave-one-out cross-validation) to distinguish between each pair of states based on their FC structure (see Methods). All pairs of conditions and/or recording sessions that presented a ground-truth accuracy lower than the 95th percentile of the accuracy obtained on shuffled datasets were not considered for further analysis.

**Figure 8.**
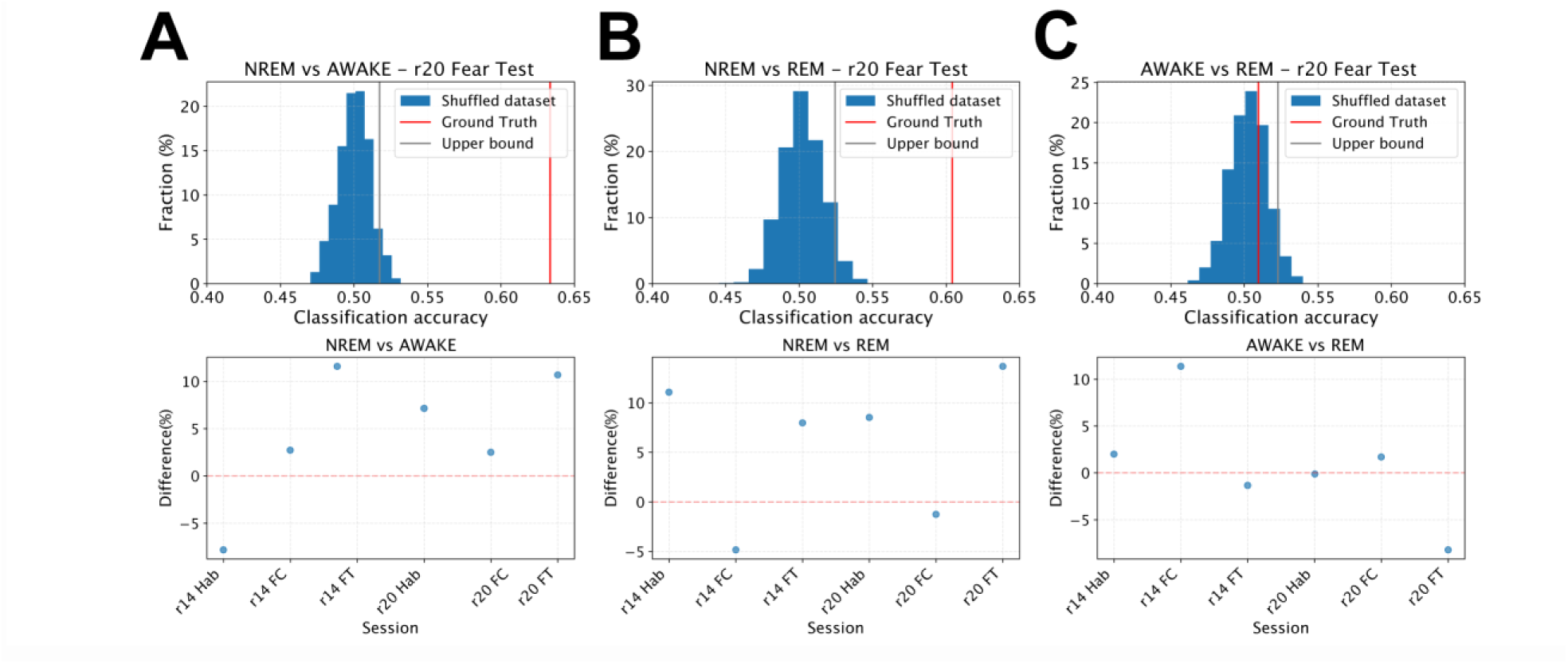
Overall classification accuracy across sessions. **A, B** and **C** (top) represent (from left to right) the histogram of classification accuracy for the shuffled datasets for NREM vs wakefulness (AWAKE), NREM vs REM and wakefulness vs REM respectively, for an example recording session (r20 Fear Test). The red bar indicates ground truth classification accuracy, and the grey bar indicates the top 95% of the obtained shuffled accuracies. The bottom row shows, per session, the difference between the ground truth accuracy (blue dots) and the top 95% of shuffled accuracies (red dashed line). Sessions were excluded in case the ground truth accuracy was lower than the top 95% of shuffled accuracies.

Moreover, as an additional exploratory control, we expanded the original pipeline to also include a comparison between REM and wakefulness (Table 7, Fig 8C). Of note, all of the awake vs REM ground-truth values were comparable or inferior to the 95% percentile obtained with the shuffled accuracy datasets (referred also as “95% upper bound” in tables and figures), in line with our a priori assumption that REM and wakefulness show comparable FC amplitude.

**Table 7:**
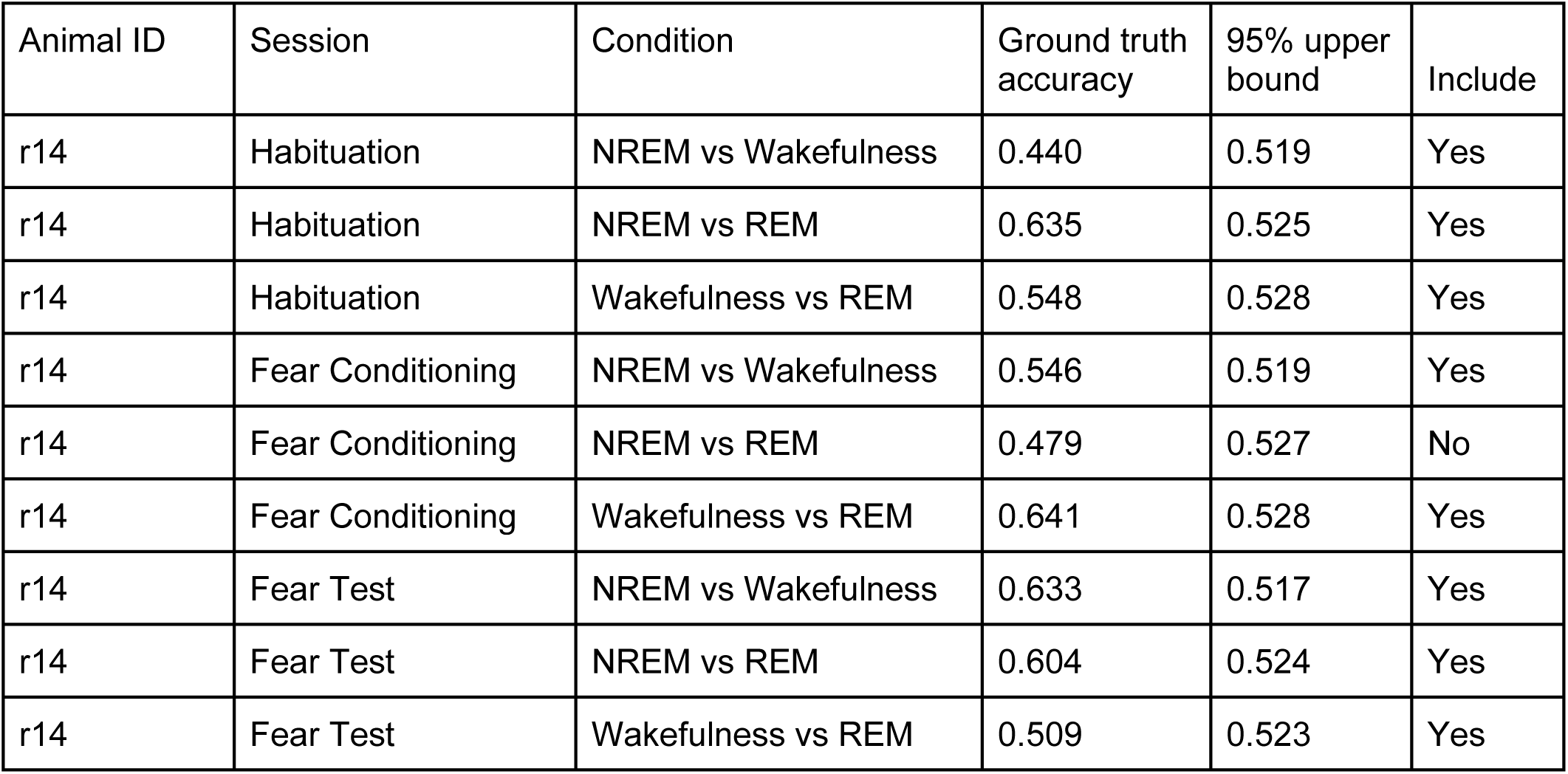
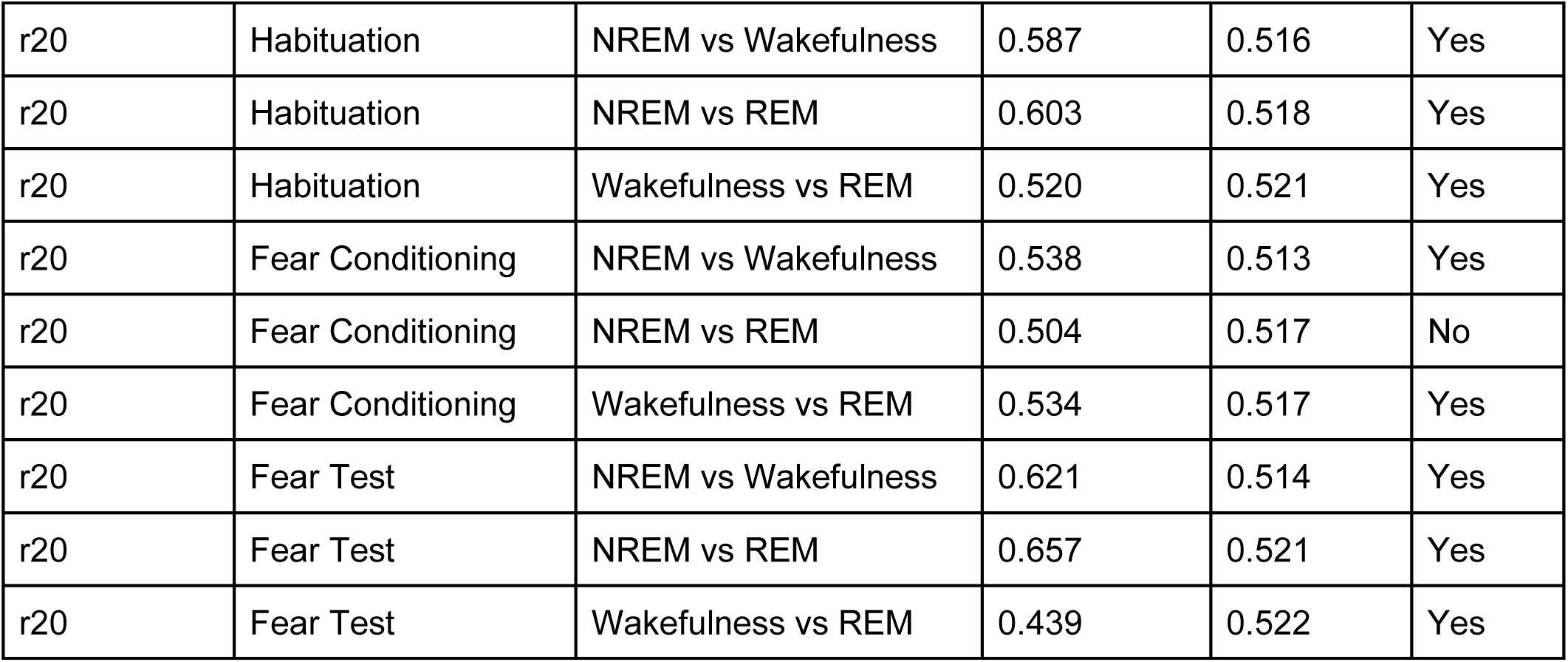
Classification accuracies between brain states across different behavioral sessions. Ground truth accuracy and 95% upper bound values are presented for each comparison. Sessions where the ground truth accuracy was lower than the top 5% percentile of shuffled accuracies were further excluded from analysis.

For some sessions, NREM vs REM could not be discriminated better than the shuffled dataset. This was in particular the case for Fear Conditioning sessions, potentially indicating that the interareal activity in the recorded regions is susceptible to different sleep dynamics depending on previous experience. Overall, we excluded two sessions based on this criterion.

#### Number of recording sessions

Finally, we assessed if the number of recording sessions was sufficiently high. In this procedure we randomized three selected sessions to compose a surrogate dataset (see Methods for details). Our results consistently indicate that we are sufficiently powered from a minimum of 4 sessions for all conditions (NREM vs wakefulness = 98.64% ± 0.006%; NREM vs REM = 96.81% ± 0.118%; wakefulness vs REM = 89.09% ± 0.028% (mean ± sem)). The minimum number of sessions to be included in the rest of the analyses was thus set to 4. Importantly, when restricted to a specific session type (Habituation, Fear Conditioning or Fear Test), our dataset was deemed to be underpowered.

Overall, our exclusion criteria indicate that, even after removing two sessions from further consideration, we had enough statistical power to draw meaningful conclusions about our hypotheses

### Functional connectivity during a fraction of NREM epochs is indistinguishable from that observed during wakefulness and REM

The first hypothesis we aimed to test (H1) asked whether a significant fraction of NREM epochs displayed FC amplitude and structure comparable to that observed in either wakefulness or REM. We first assessed data normality and then tested for significant differences between inter-areal FC values obtained between NREM and either wakefulness or REM, using Kruskal-Wallis tests with post-hoc Dunn correction (Table 8). Based on the states methods, all sessions for which no significant difference was found were not to be considered for further analysis. One recording session had therefore to be excluded since no significant differences were observed (Table 8), thus resulting in a total of 4 retained recording sessions for the NREM vs REM condition, and 6 for NREM vs Wakefulness.

**Table 8:** Normality and across conditions differences. Sessions where the ground truth p-value was higher than the 0.05 were further excluded from analysis.

| Animal ID | Session | Result of Shapiro test | Condition | Kruskal | p-value | Include |
| --- | --- | --- | --- | --- | --- | --- |
| r14 | Habituation | Not normally distributed | NREM vs Wakefulness | 0.0 | 0.0 | Yes |
| r14 | Habituation | Not normally distributed | NREM vs REM | 0.0 | 0.0 | Yes |
| r14 | Fear conditioning | Not normally distributed | NREM vs Wakefulness | 0.01 | 0.01 | Yes |
| r14 | Fear conditioning | Not normally distributed | NREM vs REM | 0.01 | 0.33 | No |
| r14 | Fear test | Not normally distributed | NREM vs Wakefulness | 0.0 | 0.0 | Yes |
| r14 | Fear test | Not normally distributed | NREM vs REM | 0.0 | 0.0 | Yes |
| r20 | Habituation | Not normally distributed | NREM vs Wakefulness | 0.0 | 0.0 | Yes |
| r20 | Habituation | Not normally distributed | NREM vs REM | 0.0 | 0.0 | Yes |
| r20 | Fear Conditioning | Not normally distributed | NREM vs Wakefulness | 0.0 | 0.0 | Yes |
| r20 | Fear Conditioning | Not normally distributed | NREM vs REM | 0.0 | 0.0 | Yes |
| r20 | Feart test | Not normally distributed | NREM vs Wakefulness | 0.0 | 0.0 | Yes |
| r20 | Feart test | Not normally distributed | NREM vs REM | 0.0 | 0.0 | Yes |

Next, we quantified the proportion of high-aiFC NREM epochs (top 25th percentile) exceeding the lowest 5th percentile of REM and wakefulness FC values (see Methods; Fig 9). Across all sessions, all high-aiFC NREM epochs exhibited higher FC values than the bottom FC values in both REM and wakefulness. Therefore, despite being generally lower, FC values in NREM sleep often overlap in magnitude to what is observed in REM and wakefulness. This observed difference is a central aspect of our hypothesis (H1), showcasing how sleep stages are heterogeneous.

**Figure 9.**
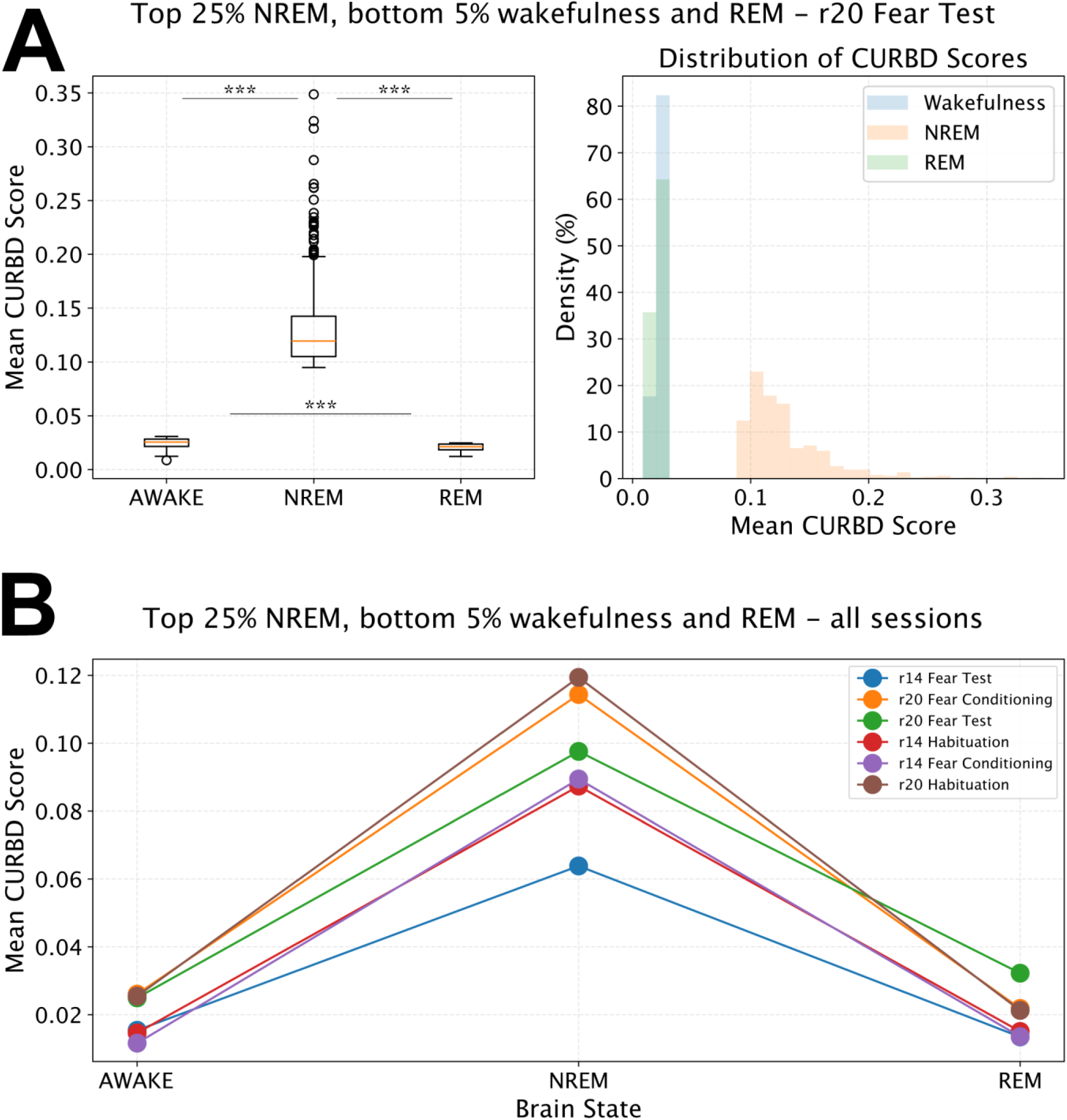
Distribution of FC magnitude in high aiFC NREM epochs and low aiFC REM and wakefulness epochs. **A.** Example Session Results: *Left*: Box plots illustrate the distribution of average inter-areal functional connectivity (aiFC) magnitude values across all sessions, per brain state. Asterisks denote a statistically significant difference (p < 0.001, Kruskal-Wallis test) for the indicated tested pairs. *Right*: Histogram displaying the overall distribution of FC magnitude. **B.** Mean FC values are presented for each session and each distinct brain state. Individual lines represent separate experimental sessions.

Lastly, we implemented a support vector machine (SVM) with leave-one-out cross-validation to test if we could discriminate high-FC NREM epochs compared to - separately - all REM and wakefulness epochs on the basis of FC structure (see Methods).

This analysis was repeated using different cutoff thresholds for FC amplitude during NREM (75th, 85th, and 95th percentiles). Based on this analysis, we observed that high-aiFC NREM epochs were indistinguishable from REM epochs at all cutoff thresholds (Table 9, Fig 10). In contrast, high-aiFC NREM epochs became indistinguishable from wakefulness epochs only when considering epochs in the top 5% of NREM aiFC values (Table 9, Fig 10).

**Figure 10.**
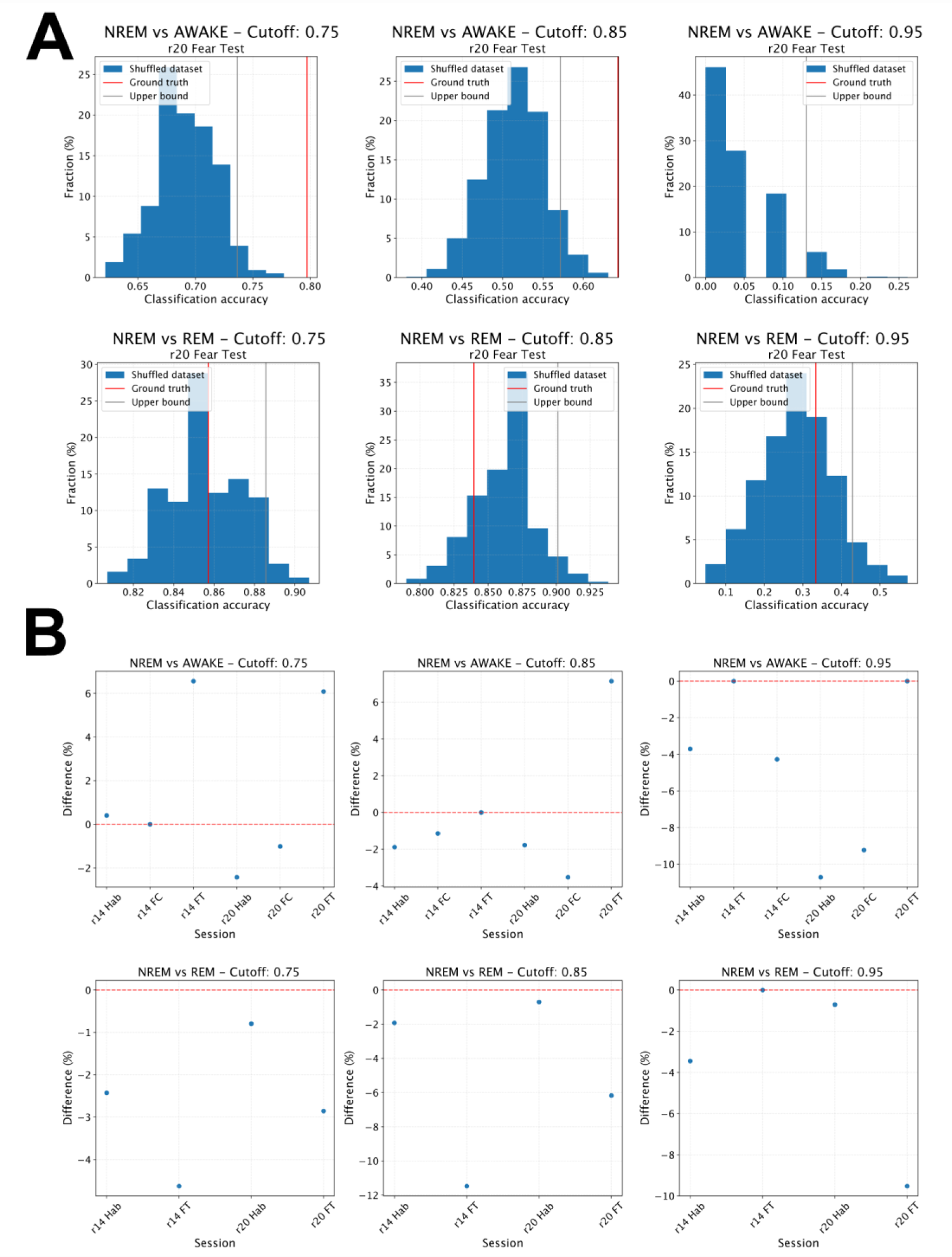
Classification accuracy across sessions for high NREM aiFC in comparison to REM and wakefulness. **A**. Results from a representative session. Top: Histograms depicting the distribution of classifier accuracy for shuffled datasets when discriminating between NREM sleep and wakefulness. Data are presented for different aiFC magnitude cutoffs used for classifier training (85th, 75th, and 95th percentiles). The red bar indicates the ground-truth accuracy, while the grey bar represents the 95th percentile of shuffled accuracies. Bottom: Same as above, but for the discrimination between NREM and REM sleep. **B.** Top: Scatter plot showing the difference between the 95th percentile of shuffled accuracies and the ground-truth accuracy for NREM vs. wakefulness classification. Each blue dot represents an individual session. The dashed red line indicates a 0% difference; sessions above this line exhibited a ground-truth accuracy exceeding the upper bound of shuffled accuracies. Bottom: Same as described above, but for NREM vs. REM sleep classification.

**Table 9:** Combined p-values across sessions for comparing NREM vs REM and NREM vs Wakefulness conditions at different statistical cutoff thresholds (75th, 85th and 95th percentile). The table shows the individual p-values from single sessions, the combined p-values using Fisher’s test, and the corresponding test statistic for each comparison and threshold combination.

| Condition | Single session P values | Magnitu<br>de cutoff<br>(percenti<br>le) | Combine<br>d p values<br>(Fisher<br>test) | Statistic<br>(Fisher<br>test) |
| --- | --- | --- | --- | --- |
| NREM vs REM | [0.69, 0.582, 0.436, 0.742] | 75th | 0.85 | 4.08 |
| NREM vs REM | [0.731, 0.634, 0.88, 0.996] | 85th | 0.99 | 1.8 |
| NREM vs REM | [0.94, 0.39, 0.806, 0.083] | 95th | 0.49 | 7.42 |
| NREM vs Wakefulness | [0.088, 0.851, 1e-16, 0.028, 0.001, 0.312] | 75th0 | 0.0 | 102.16 |
| NREM vs Wakefulness | [0.51, 0.389, 0.874, 1e-16, 0.057, 0.699, 0.566] | 85th | 0.0 | 84.77 |
| NREM vs Wakefulness | [1.0, 0.878, 0.286, 0.986, 0.077, 0.609] | 95th | 0.71 | 8.91 |

Finally, as a further control, we applied the same algorithm to classify the bottom 5% of NREM epochs versus the top 5% of REM and wakefulness epochs, under the assumption that these would represent the most distinct FC patterns between behavioral states, and should thus be effectively discriminated by a classifier. Additionally, we applied the algorithm to classify REM versus wakefulness, which is theoretically the most challenging pair to differentiate.

As expected, our algorithm successfully distinguished low NREM from high REM epochs in each recording session (Table 10). Thus, no session had to be excluded. In contrast, we were unable to reliably discriminate REM from wakefulness epochs.

**Table 10:**
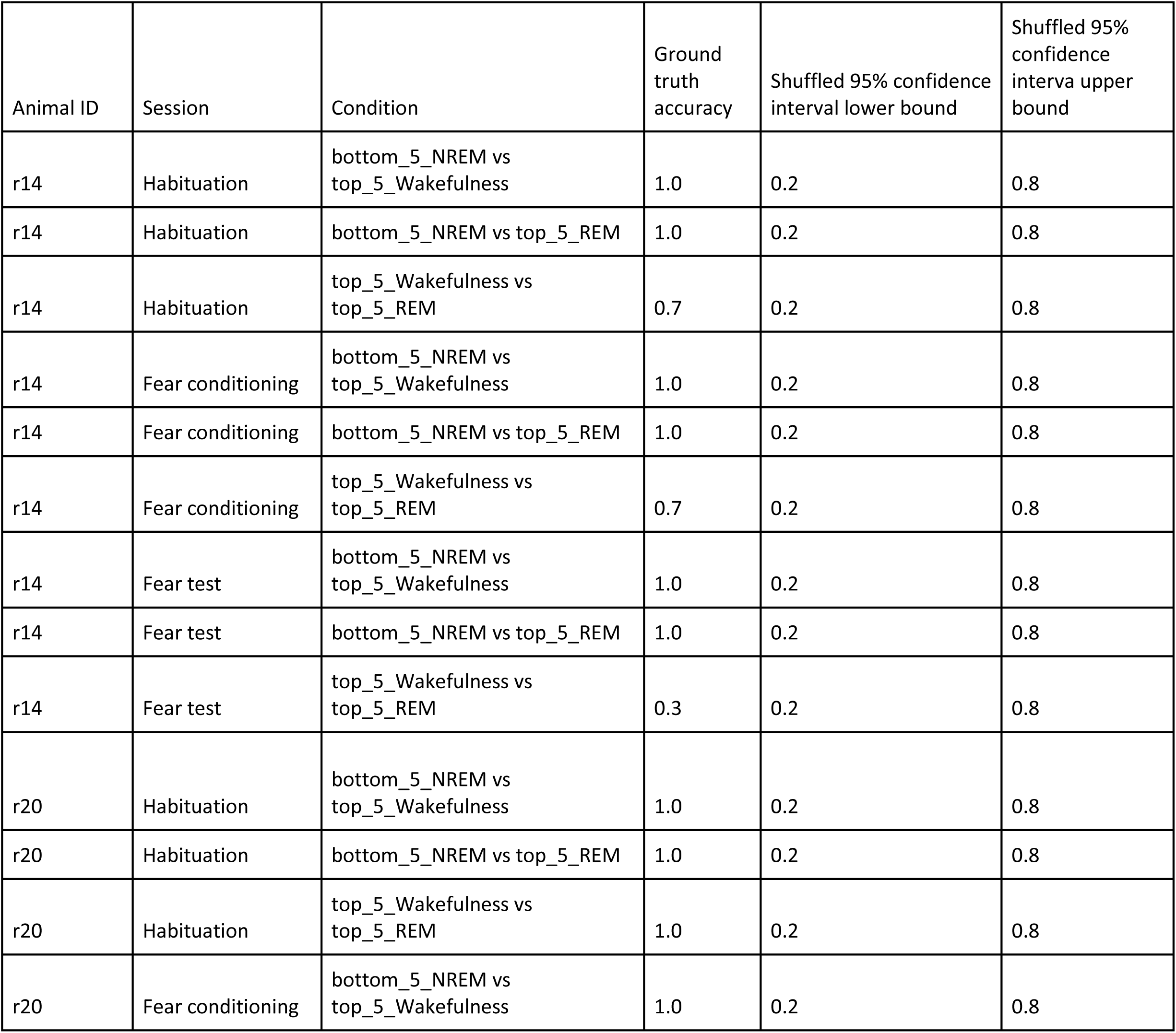
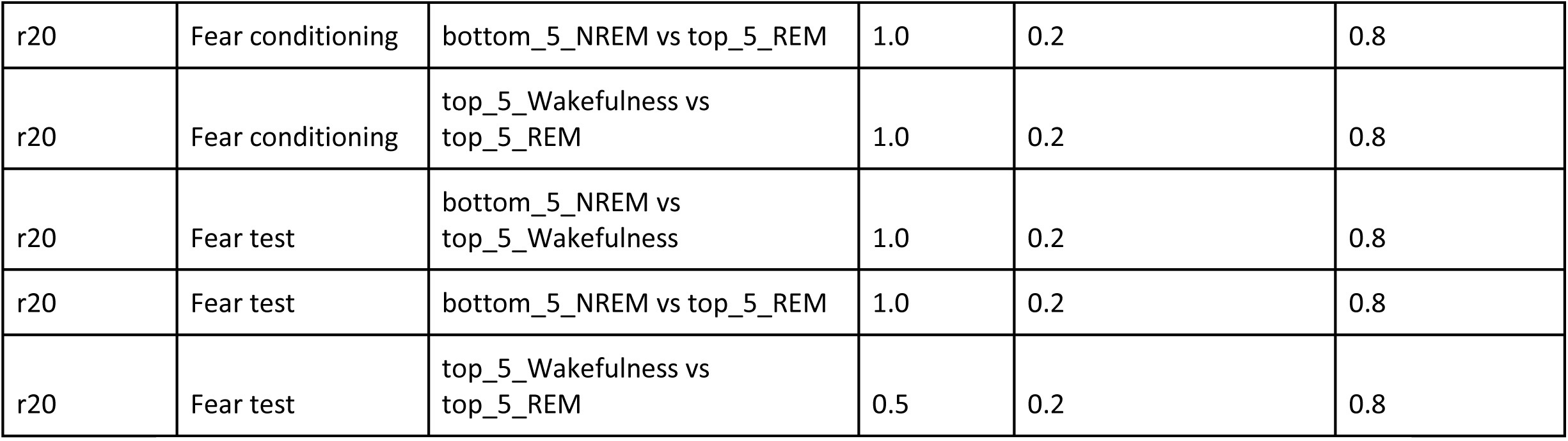
Classification accuracy across different experimental conditions for low FC NREM epochs and high FC REM and wakefulness epochs. The ground truth accuracy is shown alongside shuffled 95% confidence intervals (lower and upper bounds). The classification algorithm classifies NREM vs REM and NREM vs wakefulness successfully, as indicated by the ground truth accuracies falling outside the shuffled confidence intervals (0.2-0.8). As a control, we also applied the classifier to high FC wakefulness vs high FC REM, which the algorithm failed to resolve. Sessions where the ground truth accuracy was lower than the top 5% percentile of shuffled accuracies were further excluded from analysis

In conclusion, our results indicate that, across all examined cutoffs of FC magnitude (top 5%, top 15% and top 25%), NREM aiFC is indistinguishable from what is observed in REM sleep. This is not the case for the comparison with wakefulness, as NREM aiFC is indistinguishable from wakefulness FC only when considering epochs in NREM sleep within the top 5% range for FC magnitude. These results are in line with our hypothesis (H1), but also show that there are differences between wakefulness and REM for what pertains to which fraction of epochs shows a FC structure comparable to that observed in NREM.

### Functional connectivity during a fraction of REM epochs is indistinguishable from that observed during NREM

Following the comparison between high-aiFC NREM epochs and wakefulness/REM epochs, we proceeded to analogously investigate if REM epochs with the lowest values of FC magnitude (low-aiFC, with amplitudes below the 25th percentile) were comparable - for both amplitude and structure - to NREM epochs (H2).

First, we quantified the proportion of low-aiFC REM epochs (bottom 25%) lower than NREM epochs with the top 5% aiFC values. Across all sessions, the low-aiFC REM epochs exhibited lower FC values than the top FC epochs for NREM, indicating that, despite being generally higher, FC values in REM sleep often overlap in magnitude to what is observed in NREM (Table 11). However, it is important to note that there were differences with respect to what observed when testing H1. Here, we observed an overall lower percentage of bottom REM epochs lower than the top NREM epochs (20.01 ± 11.37 %), compared to what we found in H1 (100% of top NREM epochs were above bottom 5% REM and wakefulness).

**Table 11:**
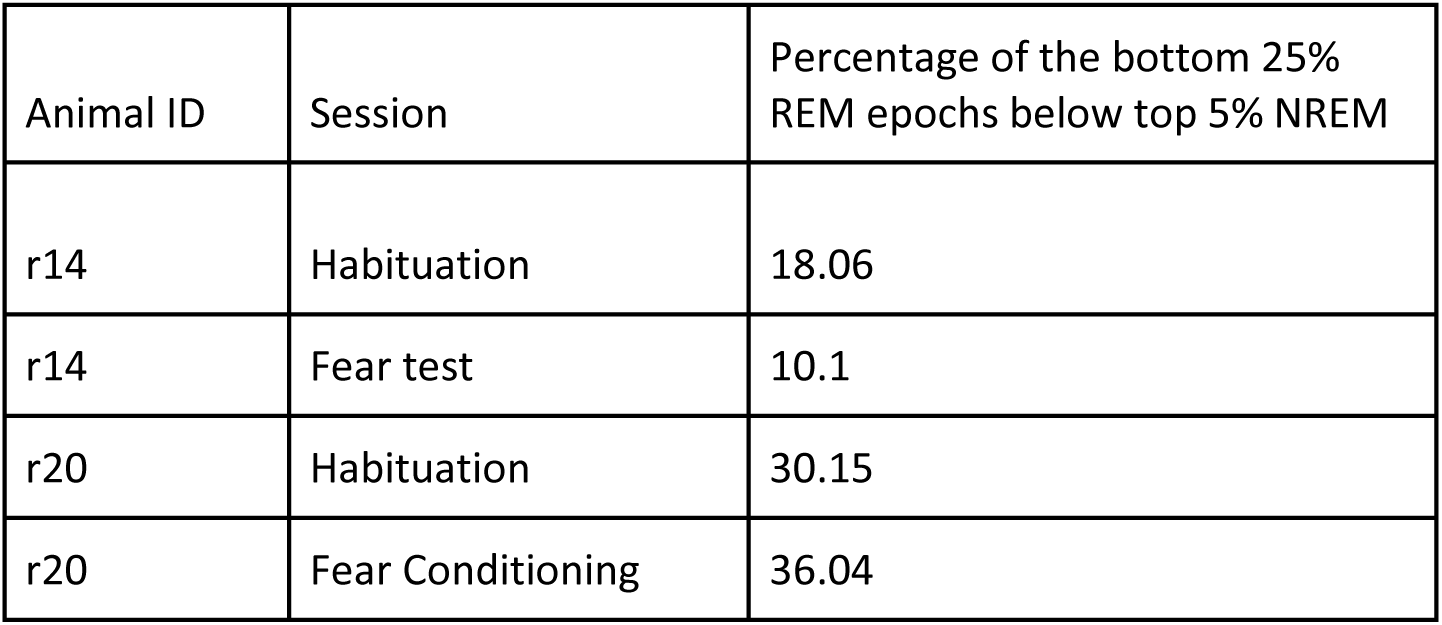
Percentage of bottom 25% REM epochs that fall below the top 5% NREM threshold across animals and experimental sessions.

Then, akin to what we performed for testing H1, we implemented a support vector machine (SVM) with leave-one-out cross-validation to test if we could discriminate low-aiFC REM epochs from NREM epochs, in terms of FC structure. This analysis was repeated using different REM aiFC cutoff thresholds (bottom 25th, 15th, and 5th percentiles). NREM and REM epochs could be discriminated when considering the bottom 25% of REM aiFC values, but not for the bottom 15% and 5% (Fig 11, Table 12).

**Figure 11.**
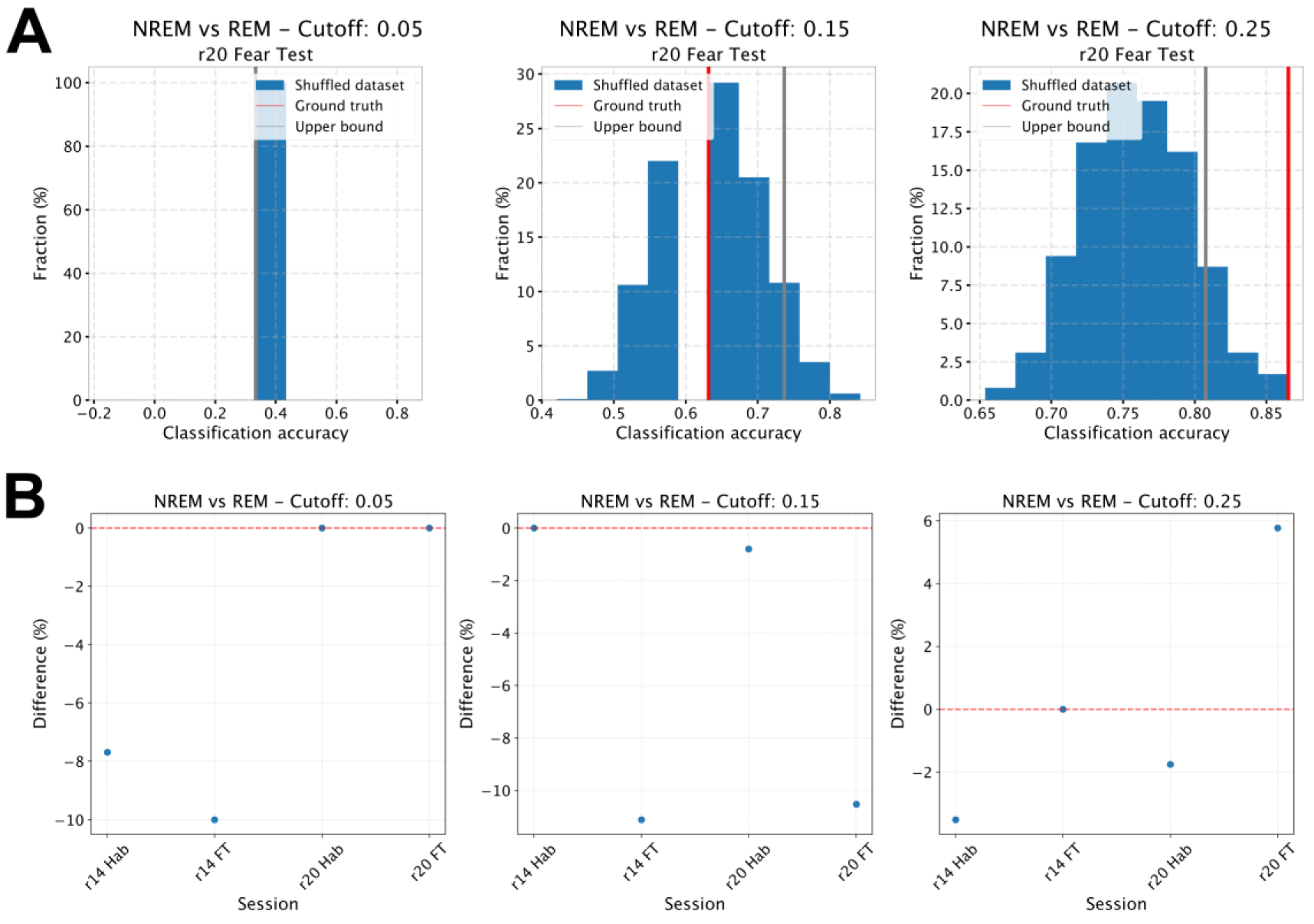
Classification accuracy across sessions for low REM aiFC in comparison to NREM. **A.** Results from a representative session. Histograms depicting the distribution of classifier accuracy for shuffled datasets when discriminating between NREM and REM sleep. Data are presented for different aiFC magnitude cutoffs used for classifier training (5th, 15th, and 25th percentiles). The red bar indicates the ground-truth accuracy, while the grey bar represents the 95th percentile of shuffled accuracies. **B.** Scatter plot showing the difference between the 95th percentile of shuffled accuracies and the ground-truth accuracy for NREM vs. REM classification. Each blue dot represents an individual session. The dashed red line indicates a 0% difference; sessions above this line exhibited a ground-truth accuracy exceeding the upper bound of shuffled accuracies.

**Table 12:**
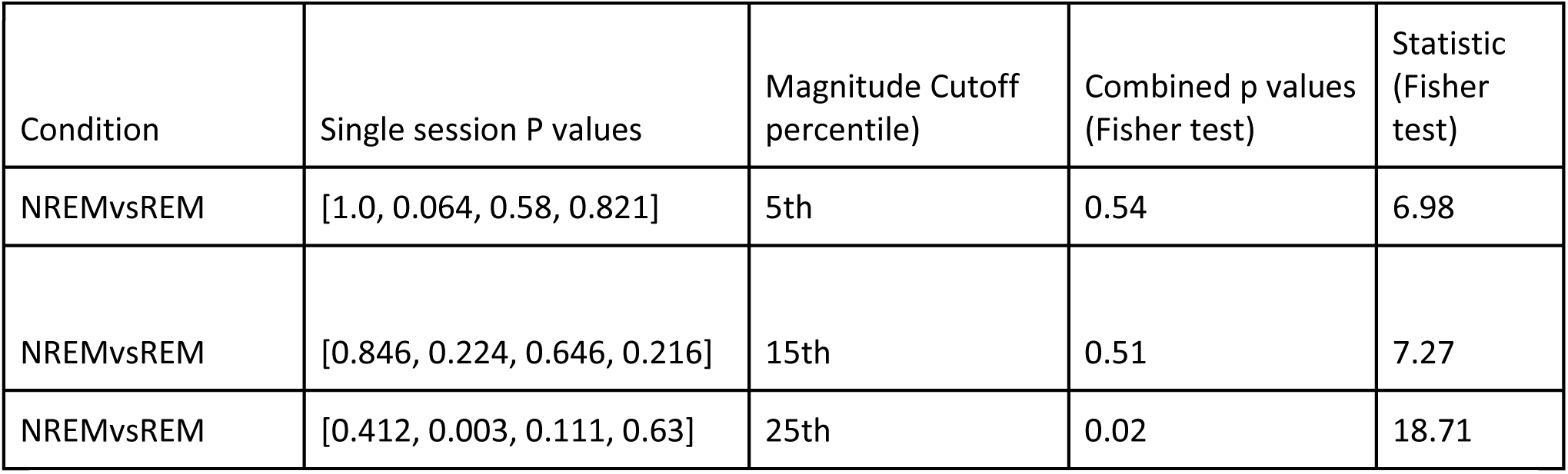
Combined p-values across sessions for comparing NREM vs REM and NREM vs Wakefulness conditions at different statistical cutoff thresholds (75th, 85th and 95th percentile). The table shows the individual p-values from single sessions, the combined p-values using Fisher’s test, and the corresponding test statistic for each comparison and threshold combination

| Condition | Single session P values | Magnitude Cutoff percentile) | Combined p values (Fisher test) | Statistic (Fisher test) |
| --- | --- | --- | --- | --- |
| NREMvsREM | [1.0, 0.064, 0.58, 0.821] | 5th | 0.54 | 6.98 |
| NREMvsREM | [0.846, 0.224, 0.646, 0.216] | 15th | 0.51 | 7.27 |
| NREMvsREM | [0.412, 0.003, 0.111, 0.63] | 25th | 0.02 | 18.71 |

Finally, as a further control, we applied the same algorithm to classify the bottom 5% of REM epochs versus the top 5% of NREM epochs. As expected, our algorithm successfully distinguished low REM from high NREM epochs in each recording session (Table 13). Therefore, no session had to be excluded.

**Table 13:** Classification accuracy across different experimental conditions for low aiFC REM epochs and high aiFC NREM epochs. The ground truth accuracy is shown alongside shuffled 95% confidence intervals (lower and upper bounds). The classification algorithm classifies NREM vs REM successfully, as indicated by the ground truth accuracies falling outside the shuffled confidence intervals (0.2-0.8). Sessions where the ground truth accuracy was lower than the top 5% percentile of shuffled accuracies were further excluded from analysis

| Animal ID | Session | Condition | Ground truth accuracy | Shuffled 95% CI lower bound | Shuffled 95% CI upper bound | Exclude session? |
| --- | --- | --- | --- | --- | --- | --- |
| r14 | Habituation | bottom_5_REM vs top_5_NREM | 1.0 | 0.2 | 0.8 | No |
| r14 | Fear conditioning | bottom_5_REM vs top_5_NREM | 1.0 | 0.2 | 0.8 | No |
| r14 | Fear test | bottom_5_REM vs top_5_NREM | 1.0 | 0.2 | 0.8 | No |
| r20 | Habituation | bottom_5_REM vs top_5_NREM | 1.0 | 0.2 | 0.8 | No |
| r20 | Fear conditioning | bottom_5_REM vs top_5_NREM | 1.0 | 0.2 | 0.8 | No |
| r20 | Fear test | bottom_25_REM vs top_5_NREM | 1.0 | 0.2 | 0.8 | No |

In conclusion, our results indicate that, across two of the three examined cutoffs of FC magnitude (bottom 5%, bottom 15%), REM aiFC is mostly indistinguishable from what is observed in NREM sleep. This is not the case for the comparison with the bottom 25% of REM sleep epochs, that we could successfully distinguish from NREM. Thus, similarly to what we observed for H1, the results that we obtained support our hypothesis that a fraction of REM epochs with low amplitude shows an FC structure comparable to that observed in NREM (H2).

## Discussion

In this study we investigated how inter-areal functional connectivity (FC) varies across brain states, in terms of both strength and structure. We observed that, in line with established findings, NREM sleep is characterized by generally weaker FC compared to both wakefulness and REM sleep (Casali et al., 2013; Massimini et al., 2005; Olcese et al., 2016; Storm et al., 2017). However, in agreement with our hypotheses, a significant fraction of epochs displays FC amplitude and structure compatible with those of a brain state other than that indicated by the ongoing neural activity patterns. Specifically, the amplitude of inter-areal FC during a significantly higher than chance fraction of NREM sleep epochs shows values as high as those typically observed in wakefulness and NREM sleep. Furthermore, the FC structure in these NREM epochs - i.e., which specific neurons are coupled to each other - is also indistinguishable from that observed in wakefulness and REM sleep. We observed analogous results for REM sleep, which displayed a significant fraction of epochs whose FC amplitude and structure was indistinguishable from what was typically seen in NREM sleep.

Interactions between neurons, here quantified in terms of FC, have been suggested by various theories to underlie the ability of the brain to generate consciousness (Koch et al., 2016; Seth & Bayne, 2022; Storm et al., 2024), with high and complex FC - such as that observed in the thalamocortical system during wakefulness and REM - indicating the presence of consciousness. In contrast, weak, simple (e.g., modular) FC is found in brain structures such as the cerebellum and in the thalamocortical system during NREM sleep. However, growing evidence suggests that brain states are not monolithic. For example, dream reports have been observed during NREM sleep (and are sometimes absent during REM sleep), but the underlying neural correlates remain controversial (Siclari et al., 2017; Wong et al., 2020). Our results suggest that epochs characterised by similar patterns of neural activity (which are commonly used to distinguish brain states) can differ in terms of FC structure. This, in turn, might support different levels of consciousness. For example, a period of NREM sleep characterized by a REM-like FC structure might correspond to a period in which dreaming occurs - and vice versa for a period of REM sleep with a NREM-like FC structure (Tononi et al., 2024).

Interestingly, our analysis suggests that the occurrence of epochs with similar FC structure is more frequent between NREM and REM than between NREM and wakefulness. This is highlighted by the observation that NREM epochs with an FC structure indistinguishable from wakefulness are only observed when NREM FC amplitude is in the top 5th percentile; conversely, a comparable FC structure between NREM and REM sleep can be observed for NREM epochs with FC amplitude in the top 25th percentile (and in the bottom 15th percentile for what pertains REM epochs). The presence of REM-like FC structure during NREM sleep (and vice versa for REM sleep) might provide a neural substrate for the reported dream phenomenology; on the other hand, the observation of limited wake-like FC structure in NREM suggests that, in line with phenomenological evidence, wake intrusion in NREM sleep is not very prevalent (Nobili, 2012; Vyazovskiy et al., 2011). An interesting aspect to pursue in future studies will be to characterize the similarity in FC structure between wakefulness, REM and the different stages of NREM sleep; this remains however challenging in rodent experiments, for which no staging of NREM sleep is commonly done (Rayan et al., 2024).

Our results, furthermore, align with recent findings suggesting that brain states such as wakefulness and sleep are more fragmented than previously thought (Olcese, Lohuis, et al., 2018; Parks et al., 2024; Vyazovskiy et al., 2011), with variations occurring at the sub-second time scale. Indeed, the picture that has emerged in recent years suggests that macro-states such as wakefulness, NREM and REM sleep are further structured into a series of micro-states that are associated with specific activity patterns as well as behavioral indicators such as pupil size fluctuations (Chang et al., 2025; Parks et al., 2024). Therefore, the different types of FC structure might represent another indication of the heterogeneous nature of brain states. This further expands previous findings indicating that coupling between different areas - and even between different sets of neurons - is differentially modulated across the wake-sleep cycle: while most pathways are depressed during NREM sleep compared to wakefulness, some are preserved or even enhanced (Olcese, Bos, et al., 2018; Olcese et al., 2016; Storm et al., 2017). Our study adds a temporal dimension to these observations and paints a picture of a very dynamic functional architecture, whose function might include well-known sleep-related phenomena such as memory consolidation (Klinzing et al., 2019), but also have relevant implications for consciousness. In particular, specific brain networks might differentially - and independently - generate conscious processing over the course of a night’s sleep, with different networks supporting for example specific forms of dreams (Nir & Tononi, 2010; Siclari et al., 2017). A similar process, furthermore, might take place during other states in which consciousness is lost, such as the unresponsive wakefulness syndrome (AWS). Many studies have now reported that a sizable fraction of AWS patients can display signs of consciousness, but that their ability - for instance - to respond to questions varies unpredictably over time (Kazazian et al., 2024; Owen et al., 2006; Van Erp et al., 2015). Such fluctuations, in view of our results, might be due to variability not so much in neural activity patterns, but rather in the FC structure.

Our study, nevertheless, suffers from some limitations. First of all, the available dataset was limited in terms of animals and experimental sessions, and the application of exclusion criteria further reduced the dataset size. This prevented us from investigating whether factors such as the contextual manipulations that were part of the original experimental paradigm had an influence on the results. Interestingly, we had to exclude all sleep sessions immediately after fear conditioning took place, since in these sessions we could not differentiate the FC structure between NREM and REM sleep. Although we are unable to determine if and to what extent fear conditioning modified the sleep-dependent FC structure (but see: Pawlyk et al., 2005), this observation suggests that daily life experiences might have a profound effect on sleep-dependent (conscious) mentation, beyond the classically described REM dreaming and sleep-dependent replay of recently acquired memory traces in the cortico-hippocampal system (Euston et al., 2007; Ji & Wilson, 2007; Wilson & McNaughton, 1994). Moreover, we were only able to assess FC between a limited number of neurons recorded from four brain regions. This prevented an in-depth characterization of area-specific modulation of inter-areal FC, and might have also contributed to the relatively high overlap between FC values across brain states (see e.g. Fig 7) – something that however could also reflect low sensitivity of the CURBD approach. Recently developed methods such as multi-area Neuropixels probe recordings (van Daal et al., 2021) make it now possible to overcome limitations in the number of simultaneously recorded neurons, and offer a promising avenue for future studies. Finally, we performed experiments in rats, a species with a different sleep architecture compared to humans. Therefore, we cannot exclude that the results that we observed can generalize to human subjects.

In conclusion, in this study we provided for the first time evidence that, while functional connectivity varies between brain states (wakefulness, NREM and REM sleep) in terms of both amplitude and structure, a sizable fraction of epochs within a given state better aligns with other brain states, in spite of the presence of neural activity patterns typical of the original state. In view of the tight connections between FC patterns and consciousness, this suggests that the level of consciousness might not just fluctuate between wakefulness, NREM and REM sleep, but also within such states, thus providing a potential substrate to explain - for example - the presence or absence of dream-like activity.

## Acknowledgments

This study was supported by the Interdisciplinary Doctorate Agreement Fund of the University of Amsterdam (to UO and LT), by an Amsterdam Brain and Cognition Support Grant (to UO and LT) and by the “Funding Consciousness Research with Registered Reports” program of the Center for Open Science (to UO). We thank Otto Márton for his support with preliminary analyses; we thank the workshop of the Universiteit van Amsterdam and Gerjan Huis in ‘t Veld for all the support.

## Data availability

In adherence to open science principles and in agreement with the journal’s policies, all data necessary to reproduce the results presented here is openly available in a dedicated repository: https://github.com/joaohnp/FC_Consciousness.

## Code availability

In compliance with the requirements for Registered Reports and in accordance with the principles of open science, all code associated with this manuscript is openly available at https://github.com/joaohnp/FC_Consciousness.

## Author Contributions

Conceptualization: JP, UO; Data curation: JP; Formal analysis: JP; Funding acquisition: UO, LT; Investigation: JP; Methodology: UO, JP, GM, JM; Project administration: UO, JP; Resources: UO, LT; Software: JP, GM; Supervision: UO; Validation: UO, LT, JM; Visualization: JP, UO; Writing – original draft: JP, UO, GM, JM; Writing – review & editing: JP, UO;

## Notes

https://github.com/joaohnp/FC_Consciousness

### Competing Interest Statement

The authors have declared no competing interest.

